# Differential contribution of HCN1 and HCN4 to the synchronisation of sinoatrial pacemaker cells

**DOI:** 10.64898/2025.12.16.694405

**Authors:** Konstantin Hennis, Chiara Piantoni, Martina Orabona, Linh Pham, Michael Schänzler, Nicolas Auerbach, Michael Hörzing, Yakun Wu, René Rötzer, Andra Kohlrautz, Colin Feldmann, Mariano Gonzalez Pisfil, Daniela Kruck, Julia Rilling, Beate Averbeck, Viacheslav O. Nikolaev, Martin Biel, Christian Leibold, Stefanie Fenske, Christian Wahl-Schott

## Abstract

The heart’s ability to beat with high precision, with a regular and steady rhythm relies on the synchronised activity of pacemaker cells in the sinoatrial node (SAN), which communicate with one another through gap junctions. This process ensures that electrical impulses are organised and reach a critical mass to ignite the electrical activity of the surrounding atrial tissue and trigger the regular heartbeat.

**Hypothesis:** We hypothesise that hyperpolarisation-activated, cyclic nucleotide-gated (HCN) channels play a pivotal role in maintaining this synchronisation process. This idea aligns with the well-established role of HCN channels in stabilising the membrane potential in the voltage range of the slow diastolic depolarisation, counteracting voltage fluctuations and effectively filtering out variations in the beating rate from neighbouring cells.

**Aim:** We focus on two specific HCN channel subtypes—HCN1 and HCN4—and their contributions to the rapid synchronisation of pacemaker cells, a phenomenon known as phasic entrainment. Using two HCN channel-mutant mouse models, we dissect the distinct roles of HCN1 and HCN4 in this process.

**Methods:** We employed patch-clamp electrophysiology to determine phase response curves (PRCs) to predict the ability of single cells to interact in the SAN. Using computer simulations, the behaviour of the SAN at the network level was determined.

**Results:** We found that HCN1, but not HCN4, is essential for the fast synchronisation of pacemaker cells in the SAN and propose a mechanism by which HCN1 channels regulate this process.

**Conclusion:** These findings highlight HCN1 as a critical component for ensuring the precise and rapid coordination needed for synchronisation of the SAN, for a regular and consistent heartbeat.

**Translational perspective:** Our work provides essential insights into the cellular and molecular mechanisms governing SAN function and lays the groundwork for several clinically relevant applications. Understanding how I_f_ blockers may affect heart rate and rhythm stability is crucial for assessing potential side effects of current and future subtype-specific HCN channel inhibitors. Moreover, our findings could support new diagnostic strategies for detecting patients at risk of SAN dysfunction. Ultimately, these findings pave the way for innovative therapeutic approaches, including targeted channel modulation and future cell or gene therapies to restore pacemaker stability.

## 1. Introduction

The regularity of the heartbeat is a key characteristic of cardiac activity, detectable by feeling the pulse. Normally, at rest, the heartbeat is almost perfectly regular, with physiological baseline fluctuations known as heart rate variability and with respiratory sinus arrhythmia causing minor changes.^1^ These natural fluctuations are attributable to a balanced influence of the autonomic nervous system. Even physical activity only induces smooth changes in heart rate (HR), without abrupt frequency changes. On the other side, in pathological conditions such as sinoatrial node dysfunction (SND) and potentially fatal secondary atrial and ventricular arrhythmias, the HR can be severely disturbed and become highly irregular.^2–5^ These examples underscore the vital importance of maintaining a regular heartbeat. The heartbeat arises in the sinoatrial node (SAN) network by spontaneous activity of pacemaker cells. In these cells, spontaneous firing of action potentials (also called pacemaker potentials) is driven by the slow diastolic depolarisation (SDD). This phase of the pacemaker cycle is initiated after the action potential when the membrane potential is most negative (maximum diastolic potential, MDP). During SDD, net inward currents slowly depolarise the membrane until the threshold potential is reached, at which the upstroke of the next action potential is triggered. These currents are generated by numerous ion channels and transporters in the cell membrane (termed *Membrane Clock*) which are functionally coupled to intracellular Ca^2+^ dynamics (termed *Calcium Clock*)^6, 7^. The rate of membrane depolarisation during SDD controls the firing frequency of pacemaker cells, determining the HR.^6, 8–11^

Hyperpolarisation-activated cyclic nucleotide-gated (HCN) cation channels generate the cardiac pacemaker current I_f_ and are crucial for generating SDD. There are four subtypes (HCN1-HCN4) in humans, mice, and other mammals. HCN4, HCN1, and HCN2 are expressed in the SAN of humans and mice, ^12–15^ with HCN4 being the predominant isoform, i.e., comprising 60-80% of sinoatrial I_f_, while approximately 20-40% is mediated by HCN1^12, 16–19^. In humans, HCN1 is distributed throughout the SAN, while in mice it is mainly expressed in the SAN head and body^12, 20, 21^. HCN channels are opened by hyperpolarisation, and in addition, their activation is facilitated by the binding of cyclic adenosine monophosphate (cAMP) to a cyclic nucleotide-binding domain (CNBD) in the C-terminus of the channels. Within the physiological voltage range, HCN channel activation is enhanced by intracellular cAMP concentrations.^22^ While the isoforms HCN4 and HCN2 are highly cAMP-sensitive, HCN1 is only weakly affected by cAMP^23–27^.

Interestingly, not all pacemaker cells in the SAN have the same firing frequency. Sub-networks of pacemaker cells in different SAN regions are characterised by a broad range of differing rates, with the fastest region, the leading pacemaker region, driving the entire SAN.^28–33^ As HR changes, the anatomical position of the leading pacemaker region shifts, so that different sub-networks of pacemaker cells are responsible for generating different HRs. Within these sub-networks, individual pacemaker cells synchronise their firing rates to a common rhythm and interact with other firing or nonfiring cell types of the network, such as cardiac fibroblasts and atrial cardiomyocytes, to reach a critical mass to drive the SAN at a stable rate and finally generate a rhythmic heartbeat. This synchronisation process, termed intrinsic entrainment, is mediated by electrical coupling via gap junctions.^34^ It is based on the principle that regular pacemaking is achieved by reducing the electrical heterogeneity present at the single-cell level. In addition to intrinsic entrainment, subthreshold fluctuations in the membrane potential of nonfiring cells could be amplified across different cells via gap junctions, eventually reaching the threshold for firing and thereby contributing to rhythmic pacemaking–a mechanism termed stochastic resonance^35–38^. Importantly, in both cases, many different components of the *Membrane* and *Calcium Clocks* may contribute to intercellular interactions, including HCN channels.

There are two types of intrinsic entrainment that take place on different time scales (*Figure 1*). The first, tonic entrainment,^20^ involves interactions between firing cells and recently discovered nonfiring pacemaker cells (*Figure 1A*; right part and *Figure 1C*).^6, 8, 13, 20, 39^ These interactions happen on a slow time-scale, i.e., within periods of 5-60 s corresponding to the duration of nonfiring episodes observed in SAN cells^20^. The second, phasic entrainment, occurs between two firing cells (*Figure 1A*; left part and *Figure 1B*) and rapidly occurs from beat to beat.^34, 39–41^ The brief interactions are caused by slight differences of a few milliseconds in the firing periods of neighbouring pacemaker cells. A fast-firing cell that is coupled to a slow-firing cell via gap junctions prematurely stimulates the slower cell late in its cycle and induces a phase advance of the next pacemaker potential (*Figure 1B*). Consequently, the cycle length is shortened, and the firing rate of the slow cell is accelerated (phase advanced). In return, the phase advanced pacemaker potential now prematurely stimulates the fast cell early in its next cycle, inducing a phase delay, lengthening the cycle, and slowing the fast cell.^34^ These mutual phasic interactions synchronise the firing rates of pacemaker cells from beat to beat within the SAN sub-networks. Both the tonic and the phasic components of intrinsic entrainment are physiologically important because together they set the natural frequency of the SAN network and stabilise the current HR set point.

**Figure 1.**
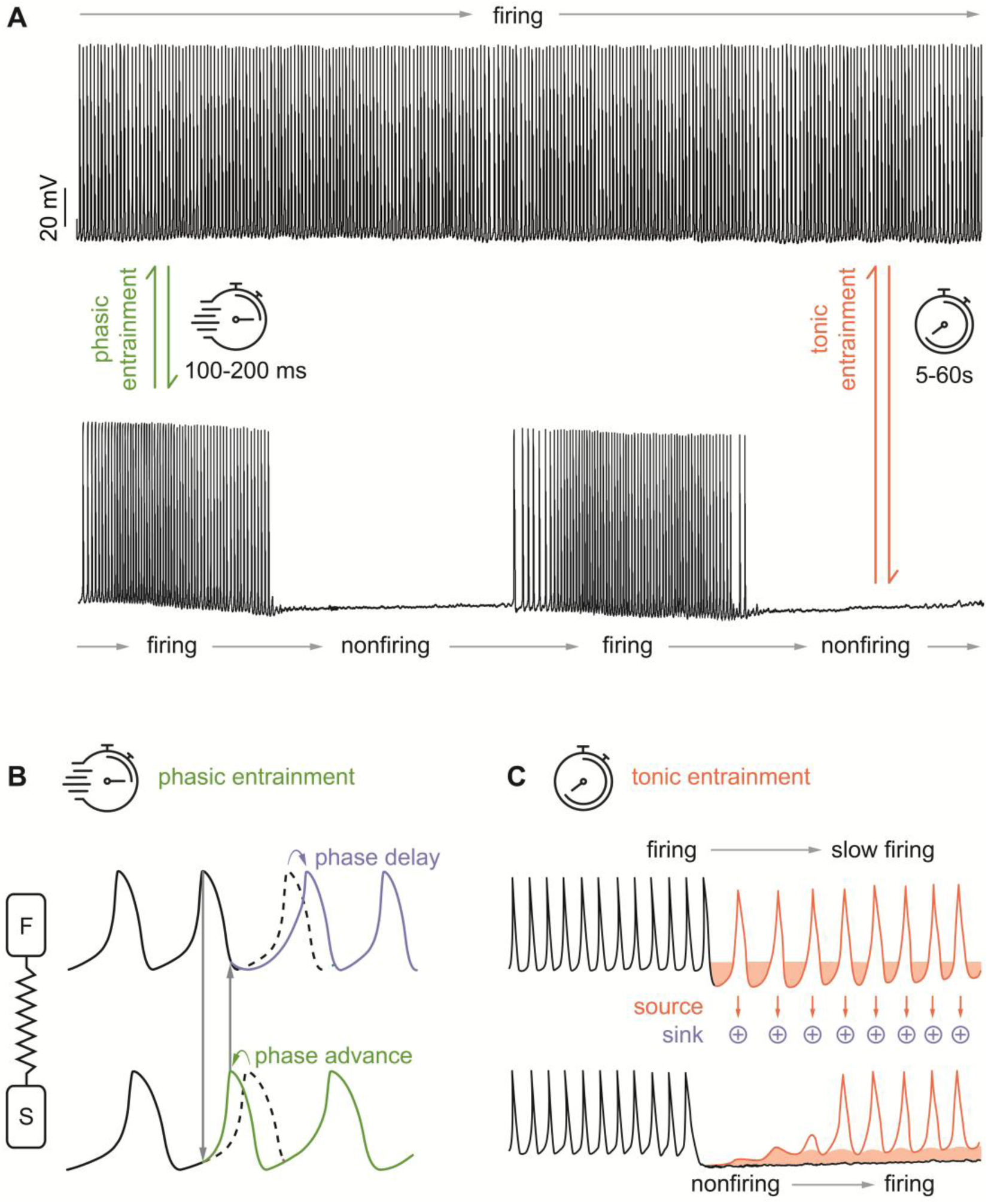
Intrinsic entrainment of pacemaker cells. *(A)* Sinoatrial pacemaker cells can either remain continuously in firing mode (upper panel) or switch between firing and nonfiring (lower panel). *(B)* Phasic entrainment between two firing pacemaker cells is based on spontaneous pacemaker potentials of a fast-firing (F) and slow-firing (S) cell connected by high-resistance gap junctions. All-or-none action potentials are conducted through gap junctions but the high coupling resistance attenuates them, so they reach the acceptor cell as subthreshold potentials, which only evoke subthreshold depolarisations in this cell. Depending on the timing in the cycle, subthreshold potentials shorten or prolong the following pacemaker potential, leading to synchronisation of cells to a common network rhythm. The slow cell (S) fires with a longer cycle than the fast-firing cell (F). The slow cell is prematurely stimulated by the fast cell late in the cycle (phase φ = 0.8) and induces a phase advance of the next pacemaker potential. This pacemaker potential prematurely stimulates the fast cell early in the next cycle and induces a phase delay. For details, see text. *(C)* Tonic entrainment is based on long-lasting interactions between firing and nonfiring cells. During nonfiring, the membrane potential is significantly more hyperpolarised than during firing, causing cationic current flow from the more depolarised firing cell to the more hyperpolarised nonfiring cell. This interaction results in tonic inhibition, slowing down firing cells and increasing the likelihood of nonfiring cells returning to the firing mode. Altogether, these tonic interactions stabilise the SAN network rhythm.^20^

HCN channels activate in the subthreshold voltage range of SDD.^6, 9, 10, 24, 42^ In the SAN, rapidly activating HCN1 channels are present, matching the time scale of phasic entrainment. Furthermore, HCN1^-/-^ mice exhibit SND with rapidly fluctuating HR.^12^ Importantly, these HR fluctuations and pronounced beat-to-beat variability are present *in vivo* and in acutely isolated SAN preparations. In contrast, spontaneous firing of single pacemaker cells is rhythmic and regular. These findings suggest that HCN1 channels could be involved in phasic entrainment and that this function arises at the network level and not at the level of single SAN cells. Additionally, control of HCN4 by the autonomic nervous system via cAMP-dependent regulation (CDR) may also contribute to phasic entrainment. The sympathetic nervous system is known to increase cAMP and CDR of HCN4, while the parasympathetic nervous system decreases them. In this way, the CDR of HCN4 helps to set the intrinsic HR, stabilise HR transitions, and protect against SND and arrhythmia.^20^ This function of CDR of HCN4 aligns with the well-known function of HCN channels in general to stabilize the membrane potential in the subthreshold voltage range in neurons and the heart, in particular during the SDD^23, 24, 43–48^. Furthermore, computer simulations also suggested that HCN channels, together with other channels, play a role in phasic entrainment^49^. Here, we investigate the role of HCN channels in the phasic entrainment process in SAN pacemaker cells of the murine heart using wild-type (WT) mice and two mouse models, one lacking HCN1^12^ and the other one lacking cAMP-dependent regulation of HCN4 (HCN4FEA)^20^.

## 2. Methods

### Animals

HCN1-deficient (HCN1^-/-^) mice were obtained from The Jackson Laboratory (B6;129-HCN1^tm2Kndl^/J).^12^ The cAMP-insensitive HCN4FEA mice (Hcn4^tm3^(Y527F;R669E;T670A)^Biel^) were generated as described previously.^20^ Transgenic Epac1-camps mice were generated as described previously^50^. HCN1^-/-^ and HCN4FEA mice were maintained on a mixed C57BL/6N x 129/SvJ background. Epac1-camps mice were maintained on a mixed 129/SvJ x C57BL/6N x FVB/N background.

Breedings of one male and two female animals were kept under SPF conditions in a 12 h dark-light cycle environment with food and water ad libitum. Ambient temperature was 22°C and humidity 60%. At P21, offspring were separated and group-housed (2–5 animals/cage) under the same environmental conditions. Animals were kept for 2–4 months and littermates were randomly assigned to experimental groups.

All animal studies were approved by the Regierung von Oberbayern, conducted in accordance with German laws on animal experimentation, and in compliance with widely accepted ethical standards. Every effort was made to keep the number of animals at a minimum.

### Gelatine-inflated heart

The gelatine-inflated heart was prepared as described previously.^20, 51^ The mouse was anaesthetised with 5% isoflurane inhalation (CP-Pharma, Germany) and sacrificed by cervical dislocation. The chest was opened, and several incisions were made into the liver. The heart was perfused with PBS via the left ventricle until it was free from blood, followed by perfusion with warm 2% aqueous gelatine solution. Immediately, cold PBS was poured onto the gelatine-filled heart and the whole body was stored at 4 °C for 1 h to let the gelatine solidify. The heart was then carefully excised and transferred into a dish containing cold PBS to remove excess gelatine and tissue. Images were taken using a stereomicroscope (Stemi 508, Carl Zeiss AG, Germany) equipped with a colour camera (AxioCam 512 colour, Carl Zeiss AG, Germany) and ZenCore 5.3 software.

### Immunofluorescence Whole-Mount SAN

The SAN tissue was dissected from a WT mouse and fixated in 4% PFA (paraformaldehyde). After permeabilization (0.5% Triton X100, 20% DMSO in PBS) and blocking in 5% NDS (Normal Donkey Serum), the tissue was incubated with guinea-pig polyclonal HCN1 antibody (1:500; Alomone Labs, Jerusalem, Israel) and rabbit polyclonal HCN4 antibody (1:500; Alomone Labs). Alexa555-conjugated anti-guinea pig (1:500; Invitrogen, Karlsruhe, Germany) and Alexa647-conjugated anti-rabbit (1:500; Invitrogen) were used as secondary antibodies. Immunofluorescent images were taken with a Leica SP8 confocal microscope.

### Isolation and electrophysiology of murine single SAN cells

Single SAN cells were isolated from HCN1^-/-^, HCN4FEA, WT, and Epac1-camps mice as described previously.^20^ In brief, the SAN region was excised and enzymatically digested using elastase (18.4 U/ml, Merck KGaA, Germany), collagenase B (0.3 U/ml, Roche Diagnostics, Germany), and protease (1.8 U/ml, Merck KGaA, Germany). Afterwards, the tissue was stored at 4°C for a recovery time of 3 hours in modified KB medium consisting of 80 mM L-glutamic acid, 25 mM KCl, 3 mM MgCl_2_, 10 mM KH_2_PO_4_, 20 mM taurine, 10 mM HEPES, 0.5 mM EGTA, and 10 mM D-glucose (KOH pH 7.4). Subsequently, cells were mechanically separated by pipetting the digested tissue 4-8 times with modified pipette tips. After gradual re-adaptation to a physiological extracellular Ca^2+^ concentration (1.8 mM), cells were used for electrophysiological recordings. For current-clamp experiments (action potential recordings), SAN cells were continuously superfused with Tyrode’s solution containing 140 mM NaCl, 5.4 mM KCl, 1mM MgCl_2_, 1.8 mM CaCl_2_, 5 mM HEPES, and 5.5 mM D-glucose (NaOH pH 7.4) at 32°C. For voltage-clamp experiments (I_f_ recordings), cells were superfused with extracellular solution (32°C) consisting of 140 mM NaCl, 5.4 mM KCl, 1 mM MgCl_2_, 1 mM CaCl_2_, 5 mM HEPES, 5 mM D-glucose, 0.3 mM CdCl_2_, and 2 mM BaCl_2_ (NaOH pH 7.4). Pipettes were filled with intracellular solution containing: 130 mM potassium aspartate, 10 mM NaCl, 2.0 mM MgCl_2_, 2.0 mM CaCl_2_, 5.0 mM EGTA, 2.0 mM Na_2_-ATP, 0.1 mM Na_2_-GTP, and 5.0 mM creatine phosphate di(tris) salt (KOH pH 7.2). For measurements under saturating cAMP concentrations, 100 µM cAMP (Merck KGaA, Germany) was added to the pipette solution and the pH was re-adjusted (KOH pH 7.2). For perforated-patch experiments, amphotericin B (EDQM, France) was dissolved in DMSO and added to the pipette solution to obtain a final concentration of 200 μg/ml.

Action potentials (AP) of spontaneously firing SAN cells were recorded with the perforated-patch or whole-cell current-clamp technique. To determine phase response curves, depolarising current pulses (+50 pA, 20 ms) were applied every 6 seconds during periods of stationary firing frequency. For example, at a spontaneous firing rate of roughly 200 beats per minute, this would correspond to a stimulus after every 20th spontaneous cycle. Because of small physiological variations in the spontaneous firing rate, the stimulus protocol sampled phases throughout the cycle length. The protocol was run until a sufficiently high number of stimuli, ranging from 9 to 189 (an average of 48), was applied to cover the entire pacemaker cycle.

Data were recorded using a HEKA EPC10 USB patch-clamp amplifier (HEKA Elektronik, Germany) and Patchmaster v2×90.2 software. Data were analysed using Fitmaster v2×91 and ClampFit 10.2.0.12 software. In this study, the phase of the pacemaker cycle was defined from peak to peak of two successive action potentials, i.e., the time point defined as zero corresponds to the peak of the first action potential and the one defined as one corresponds to the peak of the second action potential. For every stimulation event, the preceding cycle length T_0_, timepoint of stimulation Δ, and stimulus-affected cycle length T_1_ were determined. Phase shifts (T_1_/T_0_) and timepoint of stimulation (Δ/T_0_) were calculated in relation to the previous cycle length and plotted against each other to obtain a phase response curve (PRC). Depending on the magnitude of phase shifts and the overall trajectory of the curve, PRCs were classified as type 0 or type 1 and analysed separately: PRCs with a smooth transition (negative slope) from maximum delay to maximum advance were classified as type 1, whereas PRCs with an abrupt jump (discontinuity) from delay to advance were classified as type 0. To determine averaged PRCs, data points were binned by the timepoint of stimulation (Δ/T_0_) in steps of ϕ = 0.1 (early and late phases) or ϕ = 0.05 (middle part of the curve), and the mean phase shift (T_1_/T_0_) and mean phase (ϕ = Δ/T_0_) were calculated for each bin.

### Combined patch-clamp and confocal microscopy

Combined whole-cell patch clamp and confocal ratiometric FRET imaging was performed using a Zeiss LSM 980 Airy-scan 2 superresolution microscope with a W Plan-Apochromat 20x/1.0 DIC (UV)VIS-IR water immersion objective, and a HEKA EPC10 USB patch-clamp amplifier as described above. SAN cells were isolated from transgenic mice with ubiquitous expression of the cAMP FRET sensor Epac1-camps^50^. YFP and CFP fluorescence were measured with the ROI covering the entire cell area, and FRET was continuously monitored as the ratio between emission at 517-694 nm (YFP) and emission at 454-508 nm (CFP), upon excitation at 445 nm. Images were acquired in 16 bit, bidirectional line scan mode.

Background fluorescence was subtracted from the YFP and CFP channels and signals were processed by applying a Gaussian lowpass filter with cutoff frequency 14 kHz. The FRET ratio was measured at baseline (R_0_) and after reaching a steady state (R_1_) upon intracellular application of cAMP (100 µM) via the patch pipette in the whole-cell patch clamp configuration. Changes in the FRET ratio (ΔR = R_1_-R_0_) were calculated and reported as ΔR/R_0_ after normalisation to the initial (baseline) FRET ratio.

### Whole-mount SAN calcium imaging

Intact SAN preparations of 12–16-week-old C57BL/6 mice were prepared as described previously^20^ and loaded with the Calcium indicator Cal-520, AM (Abcam) at RT for 45 min. To this end, Cal-520, AM was dissolved in DMSO (2 mM) and added to Tyrode’s solution to reach a concentration of 20 µM. After Cal-520, AM was washed out, 1 mL of blebbistatin (Merck) was applied directly to the tissues for 10 minutes at a concentration of 0.2 mg/ml and then the tissues were continuously superfused with Tyrode’s solution containing blebbistatin (0.2 mg/ml) at 32 °C. Calcium signals from the SAN were recorded using a Zeiss LSM 980 Airy-scan 2 superresolution microscope with a W Plan-Apochromat 20x/1.0 DIC (UV)VIS-IR water immersion objective. Each movie (90.32×91.28 µm) was recorded for 30 s (35.6 frames/s). An excitation wavelength of 488 nm was used and emission was collected at >500 nm. After baseline measurements, a gap junction blocker (Carbenoxolone 100 µM or 1-Heptanol 10 µM in Tyrode’s solution; Merck) was added to all the 16 preparations and tissue samples were scanned as described above. Arrhythmic events were defined as ΔCL (CL_n+1_-CL_n_) >25%. Arrhythmic tissues were defined as tissues showing ≥1 arrhythmic event during the 30 seconds recording. Analysis of calcium transients frequency, variability and arrhythmic events was performed using ClampFit 10.2.0.12 and OriginPro 2023 10.0.0.154. Signal artefacts were excluded manually.

### Simulations of Phase-Coupled Oscillators

A detailed description and validation of the computer simulations can be found in the Supplementary Material. In brief, SAN cells were modelled as phase oscillators with an unperturbed period T0 that was randomly selected from a Gaussian distribution with mean and standard deviation derived from the experimental data sets. Each SAN cell reacted to a pulse from a coupled cell via a PRC that was fitted to the experimentally obtained PRCs from Figs. 5-6. For type 0 and type 1 PRCs we used the model functions

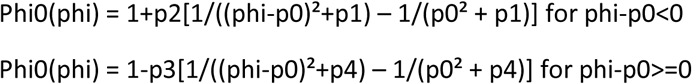

and

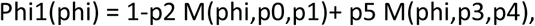

with the corrected von Mises function

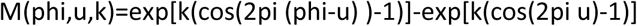

respectively. The stimulation phase phi is taken in radiants relative to the individual cycle length T0. The parameters px were fitted by minimizing Euklidian distance between model and cell averaged PRCs and are summarised in Table 1.

**Table 1.** PRC fit parameters obtained from the 6 experimental series from Figs. 5-6.

|  | Type 0<br>p0 | Type 0<br>p1 | Type 0<br>p2 | Type 0<br>p3 | Type 0<br>p4 | Type 1<br>p0 | Type 1<br>p1 | Type 1<br>p2 | Type 1<br>p3 | Type 1<br>p4 | Type 1<br>p5 |
| --- | --- | --- | --- | --- | --- | --- | --- | --- | --- | --- | --- |
| HCN1<br>WT | 0.412 | 0.0350 | 0.0197 | 0.117 | 0.174 | 0.595 | 0.576 | 1.051 | 0.559 | 0.159 | 2.00 |
| HCN1<br>KO | 0.419 | 0.0257 | 0.0100 | 0.106 | 0.159 | 0.550 | 4.98 | 0.196 | 0.134 | 0.108 | 0.963 |
| HCN4<br>WT | 0.477 | 0.0223 | 0.0100 | 0.0907 | 0.156 | 0.459 | 0.888 | 1.91 | 0.436 | 1.08 | 1.79 |
| HCN4<br>FEA | 0.470 | 0.0168 | 0.0101 | 0.0817 | 0.135 | 0.601 | 0.511 | 1.05 | 0.572 | 0.134 | 2.00 |
| cAMP<br>WT | 0.432 | 0.0288 | 0.0106 | 0.235 | 0.336 | 0.553 | 1.06 | 2.00 | 0.535 | 1.00 | 1.83 |
| cAMP<br>HCN4<br>FEA | 0.474 | 0.150 | 0.0758 | 0.0491 | 0.0889 | 0.609 | 0.907 | 1.94 | 0.597 | 0.736 | 1.85 |

For each cell, the phase of the PRC was normalised according to its individual period T0, and the effect size of the phase response was scaled by a global coupling parameter of 0.1.

Since the gap-junction coupling not only transmits a brief stimulus pulse as in the experiments but exerts a positive Ohmic current input through the whole duration of the action potential, we modelled coupling via effective PRCs that are obtained by convolving the measured PRCs with model function APM(t) of the action potential shape. For APM(t), we assume an infinitely steep rise followed by a linear decline to half amplitude in an interval of 16 % cycle length. APM(t) was normalised to area 1 before convolution.

The SAN network was modelled as a network of 500 cells randomly positioned on the nodes of a 60×12×4 cuboid, reflecting the relative geometry of the SAN with each cell coupled to the 26 next neighbours. Type 0/1 property was randomly assigned to the model SAN cells according to the fractions found in experiments.

Noise was implemented by random phase shifts. After each AP an additional random phase shift was assigned from a Gaussian distribution of mean 0 and variance dphi/SNR², where dphi was the time to the next spike estimated from the PRC and SNR = 7.3.

The network was simulated over a period of 5 min for each condition. Oscillator spike times were binned in 6.65 ms intervals resulting in a time series with a sample rate of 150 Hz. Data was low-pass filtered (30 Hz, 4^th^-order Bessel) before peak detection (peak_prominence 2). Following peak detection, tachograms and Poincaré plots were generated from the peak time series using OriginPro 2023 software (OriginLab Corporation). Tachograms were generated by plotting all consecutive cycle lengths (CL) against the time. Poincaré plots were generated by plotting each cycle length (CL+1) against the previous one (CL) to visualise beat-to-beat dispersion in cycle length.

### Statistical analysis

Statistical analyses were performed using OriginPro 2023 software (OriginLab Corporation, USA). All data are expressed as mean ± SEM. Unless otherwise stated, comparison between two groups was made using Student’s two-sample t-test. When more than one condition was applied, groups were compared using Holm’s-Sidak post-hoc test following two-way ANOVA, as indicated in the figure legends. For all statistical tests, p < 0.05 was considered significant.

## 3. Results

The present study aimed to characterise the phasic entrainment in murine SAN pacemaker cells, to determine whether this process depends on HCN channels, and if so, how HCN-dependent regulation of phasic entrainment contributes to the stability and synchronisation of the SAN. Phasic entrainment was assessed by determining phase response curves (PRCs) that quantify the phase advance and delay upon a stimulus (*Figure 1B*).^52–58^ This approach is powerful for assessing how well an individual pacemaker cell’s rhythm integrates into the network rhythm without requiring detailed information on each individual ion channel governing the pacemaker potentials.^59, 60^ In our experiments, we utilised WT mice and two previously published mouse models for SAN dysfunction.^12, 20, 61^ In the HCN4FEA mouse model, three point mutations in the intracellular C-terminus completely abolish cAMP binding to the CNBD, resulting in the expression of HCN4 channels that lack CDR (HCN4FEA).^20^ The HCN1^-/-^model lacks HCN1 channels.^12^ Given that HCN1 channels are rapidly activating and given that activation and voltage dependence of HCN4 are regulated by cAMP, we hypothesised that both HCN4 and HCN1 channels may be involved in phasic entrainment.

### 3.1. Single pacemaker cells from the sinoatrial node exhibit pronounced I_f_

Pacemaker cells were isolated using anatomical landmarks and immunostaining for the SAN marker protein HCN4 (*Figure 2*). The SAN is located in the right atrium at the border to the pectinate part of the right atrium (*Figure 2A*). The rostral part of the SAN is located close to the ostium of the superior vena cava. The caudal part extends towards the inferior vena cava, and laterally, the SAN is limited by the crista terminalis. In line with the established expression patterns of HCN1 and HCN4 in the SAN^12, 20^, immunostainings confirm the presence of HCN1 channels in pacemaker cells in the head and body region, and HCN4 channels throughout the SAN, i.e., in the head, body, and tail region (*Figure 2B*). Importantly, the homogenous distribution of HCN4-positive cells does not indicate the presence of two anatomically distinct SANs as reported for the human SAN.^62^ It also does not support the presence of separated internodal or interatrial bundles as reported for the rabbit^63^ and human heart.^64–67^ From the marked region (*Figure 2A* dotted line), SAN tissue was dissected, and individual pacemaker cells were isolated for electrophysiological recordings (*Figure 2C*).^51^ In this study, the main types of pacemaker cells, i.e., short spindle and elongated cells were investigated (*Supplementary Figure 1*). These cells showed a prominent hyperpolarisation-activated current I_f_ (n = 13, *Figures 2D* and *E*).

**Figure 2.**
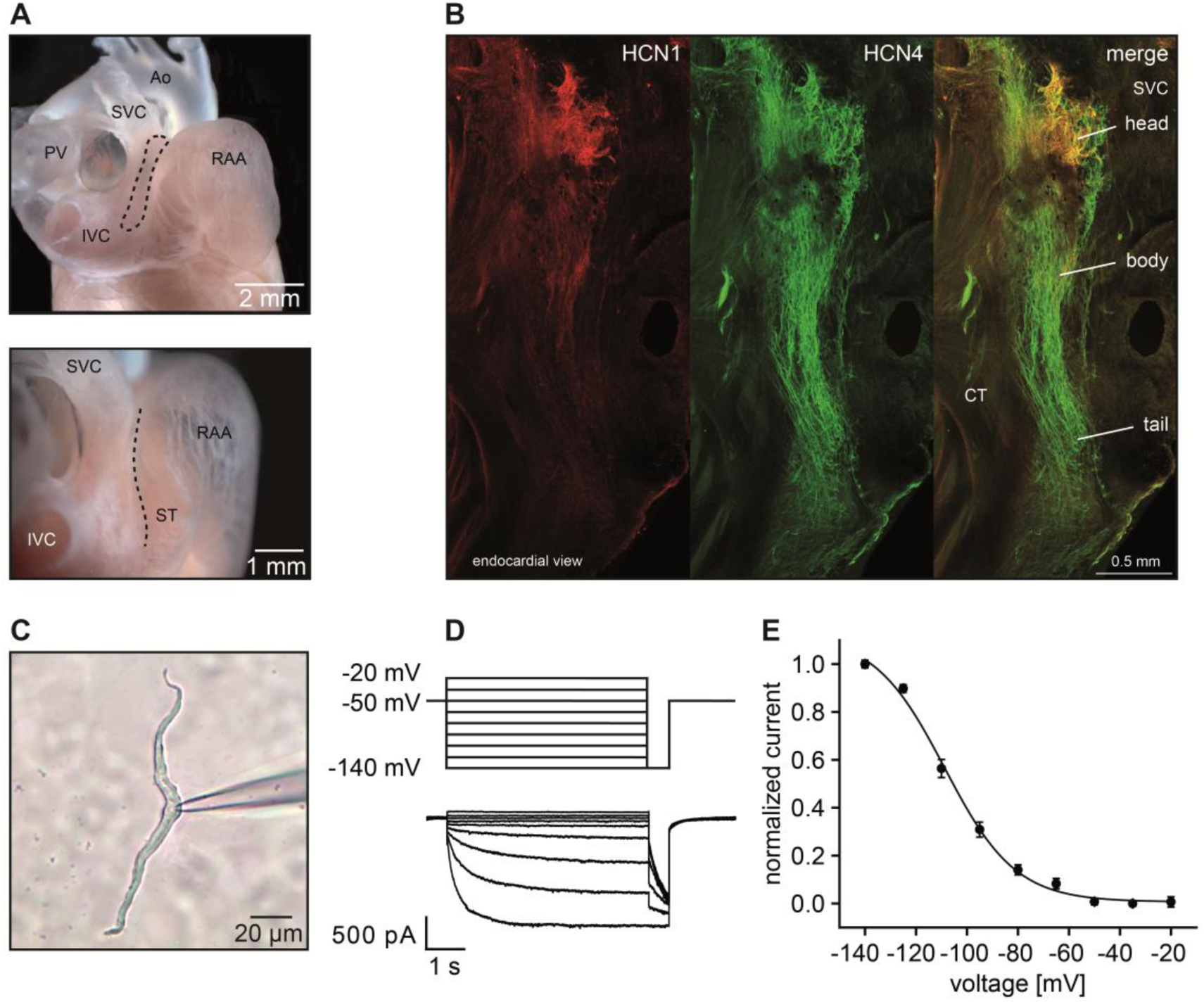
The SAN network forms the intrinsic leading pacemaker region of the heart driven by If-conducting cells. *(A)* Preparation of a gelatine-inflated heart indicating the SAN region (dashed line). *(B)* Fluorescence imaging of an anti-HCN1 (red signal) and anti-HCN4 (green signal) immunostained whole-mount right atrial preparation of a WT heart, revealing the anatomical extension (head, body, tail region) and spatial HCN1/HCN4 channel expression profile of the SAN. *(C)* Isolated SAN pacemaker cell (elongated cell) during a patch clamp recording. *(D)* Family of current traces (If) recorded from a WT pacemaker cell of the sinoatrial node using the voltage protocol shown on the top. From a holding potential of −50 mV, a series of voltage pulses ranging from −140 to - 20 mV (duration 5 s, delta V: 15 mV) were applied. *(E)* Mean activation curve of native If recorded from WT SAN pacemaker cells (n=13). Additional abbreviations: CT, crista terminalis; SVC, superior vena cava.

### 3.2. Properties of PRCs of WT pacemaker cells

PRCs were first determined in WT pacemaker cells (*Figure 3*).^34, 52–58, 68^ To this end, spontaneous pacemaker potentials were recorded using current-clamp experiments on isolated single pacemaker cells. The overall firing rate of cells included in this study was approximately 240 bpm, which is considerably lower than baseline HR observed in mice *in vivo*. This difference is well known and in line with the values reported in literature (*Supplementary Figure 2*). During periods of stationary firing frequency, brief depolarising current pulses (+50 pA, 20 ms) were applied at varying time points throughout the pacemaker cycle, and responses were recorded *(Figure 3 A* and *B*). In this study, the phase ϕ of the pacemaker cycle is defined as the period from the peak of one AP (ϕ = 0) to the peak of the next AP (ϕ = 1). From these recordings, the phase shift T_1_/T_0_ was calculated for each stimulus and plotted as a function of the timing of stimulus application within the cycle (phase ϕ = Δ/T_0_) to generate a PRC (*Figure 3C* and *D*). Responses varied with the phase of pulse application: a depolarising stimulus during the repolarisation phase prolonged the pacemaker cycle (phase delay, T_1_/T_0_ > 1; *Figure 3A and B*, upper panels), while a stimulus during the SDD phase shortened it (phase advance, T_1_/T_0_ < 1; *Figure 3A* and *B,* lower panels). The magnitude of phase shifts depends on the strength and duration of the stimulus, which were kept constant in our experiments, and on the membrane resistance.

**Figure 3.**
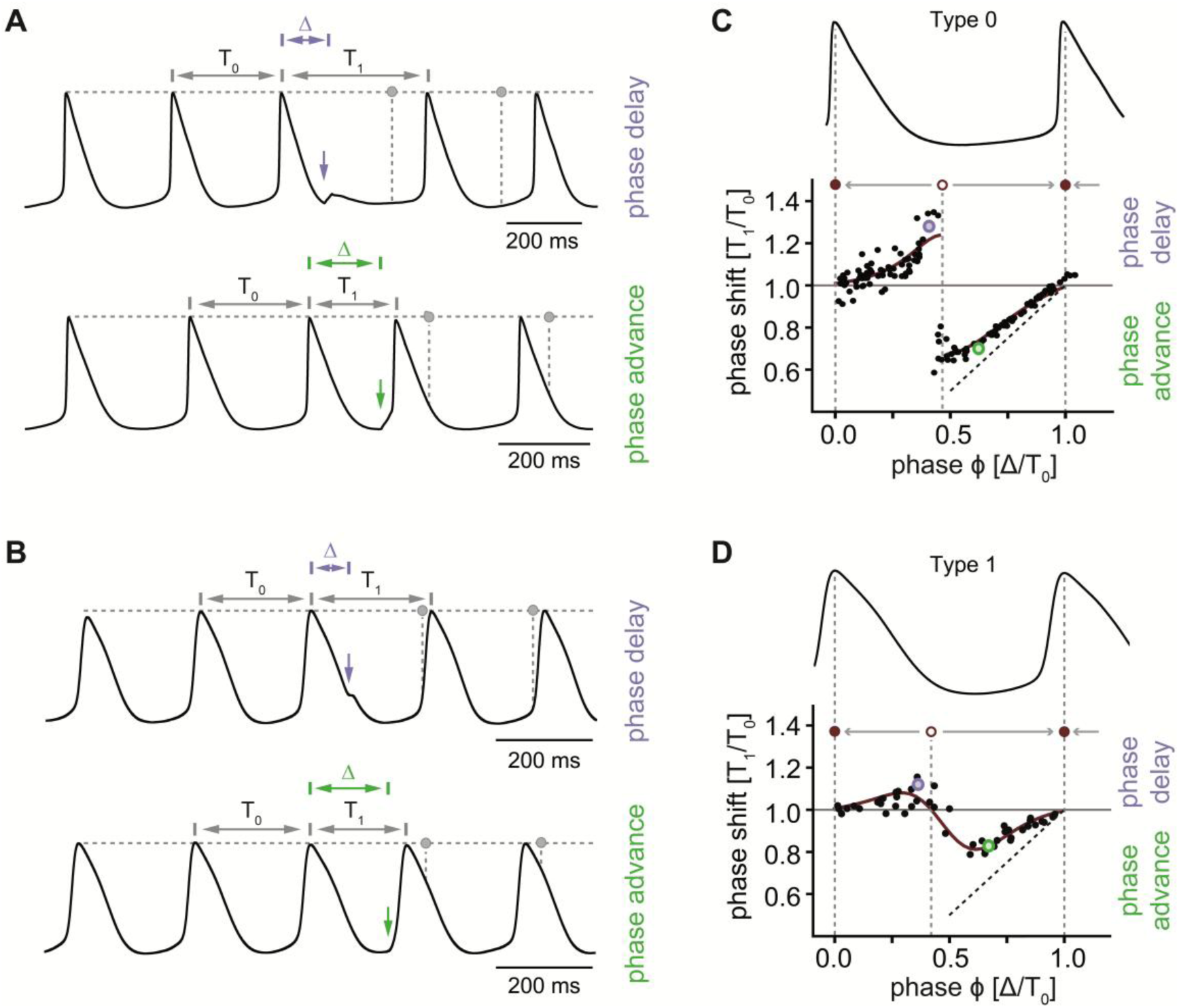
Phase response curves derived from current clamp measurements of isolated WT SAN pacemaker cells. *(A)* Protocol and representative action potential (AP) recordings for determining phase response curves. T0: spontaneous cycle length, T1: cycle length after premature stimulus, Δ: timing of premature stimulus. Please note that in this study the AP phase φ is defined from peak to peak. Brief depolarising current pulses (+50 pA, 20 ms; purple and green arrows) are applied at random time points throughout the AP phase. A depolarising impulse that occurs early in the AP phase (upper panel, ΔT/T0 = φ = 0.25) leads to prolongation of the pacemaker cycle (phase delay), whereas a late impulse (lower panel, ΔT/T0 = φ = 0.75) results in shortening of the pacemaker cycle (phase advance). The magnitude of the phase shift depends on the strength and duration of the stimulus and the membrane resistance. Grey circles mark the spontaneous rhythm of a pacemaker cell, i.e., the expected peak of an unperturbed action potential. The relatively large disturbances in panel A are reflected by the typical type 0 shape of the phase response curve shown in C. *(B)* Representative action potential recordings with milder responses to depolarising current pulses (+50 pA, 20 ms) due to a lower membrane resistance. Grey circles mark the predicted unperturbed rhythm of a pacemaker cell as in A. *(C)* Action potential trace (upper panel) and representative phase response curve (lower panel) obtained by plotting the phase shift (T1/T0) as a function of the timing of premature depolarising stimulation (phase φ = Δ/T0). The time scale of the AP trace (upper panel) matches the phase value (x-axis) in the graph (lower panel). Phase shift T1/T0 > 1 is defined as phase lengthening or phase delay. Phase shift T1/T0 < 1 is defined as phase shortening or phase advance. The 45° straight dashed line (causality line; T1/T0 = φ) indicates the theoretical upper limit for phase advance (see text for details). The relatively large disturbances shown in A are reflected by the typical type 0 shape of the phase response curve. Grey arrows point to the stable equilibrium point (filled circle, attractor). Open circle: unstable equilibrium point. Arrows point away from this point (repeller; see text for details). *(D)* Representative type 1 phase response curve caused by the relatively small disturbances shown in B.

In *Figures 3* and *5-6*, the PRCs are plotted along with a 45° straight dashed line (T_1_/T_0_ = ϕ) that represents the theoretical upper limit for phase advance. This limit is achievable only by suprathreshold stimuli that immediately trigger an action potential (*Figure 3A*, lower panel). The line is referred to as the *causality line*,^54, 69^ with *causality* implying that the response (phase advance) cannot precede its cause (premature stimulus). In practice, maximal phase advances in the PRC are close to, but not exactly on this line due to a time lag between the stimulus and the action potential peak. A large number of points near the causality line indicates maximal responses caused by suprathreshold stimulation (*Figure 3C, Figure 5D* left panel*, Figure 5H* left panel*, Figure 6D* left panel). If the points are further away from the straight line, the magnitude of phase shifts is smaller and caused by subthreshold stimulation (*Figure 3D, Figure 5D* right panel, *Figure 5H* right panel, *Figure 6D* right panel).

In WT SAN cells, two distinct types of PRCs were observed upon probe stimulation (+50 pA, 20 ms).^53,70^ Type 0 PRCs^53, 70^ are characterised by large phase shifts, with late stimulations approaching the theoretical maximum phase advance (causality line, dashed line in *Figures 3* and *5-6*), indicating close-to-threshold or suprathreshold stimulation (*Figure 3A* and *C*). In contrast, type 1 PRCs^53, 70^ show smaller shifts, suggesting that the same stimulus intensity is subthreshold in these cells (*Figure 3B* and *D*). These differences are not due to variations in series resistance or cell capacitance, as these parameters were similar in both groups (series resistance in type 0: 18.7 ± 2.6 MOhm (n = 17) and type 1: 23.9 ± 5.2 MOhm (n = 14), p = 0.37; cell capacitance: type 0: 60.1 ± 9.8 pF (n = 16) and type 1: 64.7 ± 8.4 pF (n = 11), p = 0.72). In WT pacemaker cells, more type 0 (61.1 %) than type 1 PRCs (38.9 %) were observed in the absence of cAMP (*Figure 5C*). Adding 100 μM cAMP to the intracellular solution increased type 1 PRCs to 64.3 % and reduced type 0 PRCs to 35.7 % (*Figure 5G*).

From the two types of PRCs, the maximum phase shortening, lengthening, and the entrainment range (amplitude from maximum to minimum of the PRC) were derived.^52, 60^ In WT pacemaker cells with type 1 PRCs (n = 14), the maximum phase advance was 20.0 ± 2.2 %, the maximum phase delay was 8.8 ± 2.2 %, and the entrainment range was 28.8 %. The area under the curve was 0.053 ± 0.007 for phase advance and 0.021 ± 0.005 for phase delay (*Supplementary Table 1*). There is a smooth transition (negative slope) from maximum delay to maximum advance between ∼30-60 % of the intrinsic cycle length in the middle part of type 1 PRCs (*Figure 3D*).

The PRCs are plotted with a horizontal line (T_1_/T_0_ = 1), indicating no change in cycle length (*Figures 3* and *5-6*). Type 1 PRCs intersect this line at three points: These points represent equilibrium points where phase correction in the next cycle is zero.^49, 52, 53, 58, 68, 71–73^ In WT cells, the equilibrium point at the transition from phase delay to phase advance (*Figure 3D*; open purple circle) is at ϕ = 0.42 ± 0.03 (n = 14), with a slope less than zero. This negative slope indicates an unstable equilibrium (repeller),^74^ meaning minor deviations near this point amplify, moving the rhythm towards a stable equilibrium (attractor; closed purple circles in *Figure 3C* and *D*). In contrast, the equilibrium points at the transition from phase advance to phase delay (*Figure 3C* and *D*; closed purple circle) occur at a phase of 0 and 1, with a slope greater than zero. These positive slopes indicate stable equilibrium points (attractors),^49, 52, 53, 58, 68, 71–73^ where minor phase deviations are corrected, returning the phase to the equilibrium point (attractor; at phases 1 and 0; *Figure 3C* and *D*). Depolarising stimulation early in the cycle, shortly after the stable equilibrium point, induces phase lengthening, while stimulation late in the cycle, shortly before the stable equilibrium point, induces phase shortening. This creates a negative feedback loop that stabilises the network rhythm.

In contrast, WT pacemaker cells with type 0 PRCs (n = 17; example in *Figure 3A* and *C*) exhibited a maximum phase advance of 47.8 ± 4.6 %, maximum phase delay of 36.2 ± 6.6 %, and an entrainment range of 84 %. The area under the curve was 0.126 ± 0.01 for phase advance and 0.059 ± 0.008 for phase delay (*Supplementary Table 1*). These PRCs showed a steep positive slope of phase delay and phase advance, with stable equilibrium points at 0 and 1, and an abrupt jump (discontinuity) from delay to advance after ∼40 % of the intrinsic cycle length. At this discontinuity, phase resetting is highly sensitive to the stimulus phase.

### 3.3. cAMP-dependent regulation of HCN4 does not contribute to phasic entrainment

To assess if phasic entrainment depends on CDR of HCN4, PRCs were determined from SAN cells of WT and HCN4FEA mice. In WT, HCN4 channels are regulated by intracellular cAMP, while in HCN4FEA mice, CDR is absent.^20^

To first test whether intracellular application of cAMP via the pipette solution is possible and effective, we combined confocal ratiometric FRET imaging using a Zeiss 980 Airy-scan 2 super-resolution laser scanning microscope with whole-cell patch clamp (*Figure 4*). To this end, we isolated SAN cells from transgenic mice with ubiquitous expression of the cAMP FRET sensor Epac1-camps (*Figure 4A* and *B*)^50^. Upon binding of cAMP to the FRET sensor, a conformational change leads to a spatial separation of CFP and YFP, thereby to a decrease in FRET, characterized by a concomitant decrease in YFP and increase in CFP fluorescence (*Figure 4A*). After gigaseal formation and subsequent rupture of the membrane patch, i.e., establishing the whole-cell configuration, cAMP (100 µM) entered the cell from the pipette solution, leading to a decrease in the FRET ratio (ΔR/R_0_) of 10.7 ± 1.7 % (n = 11) (*Figure 4C*). This demonstrates effective intracellular application and global and homogenous distribution of cAMP throughout the cell (*Supplementary Video 1*) within 15.6 ± 2.7 seconds (n = 11), as determined by measuring the time from rupture of the membrane patch to reaching a new steady state in the FRET ratio (R_1_). Importantly, the experiments indicate that, at the cAMP concentrations used, all regions of the SAN cells–including potential microdomains–are accessible to cAMP.

**Figure 4.**
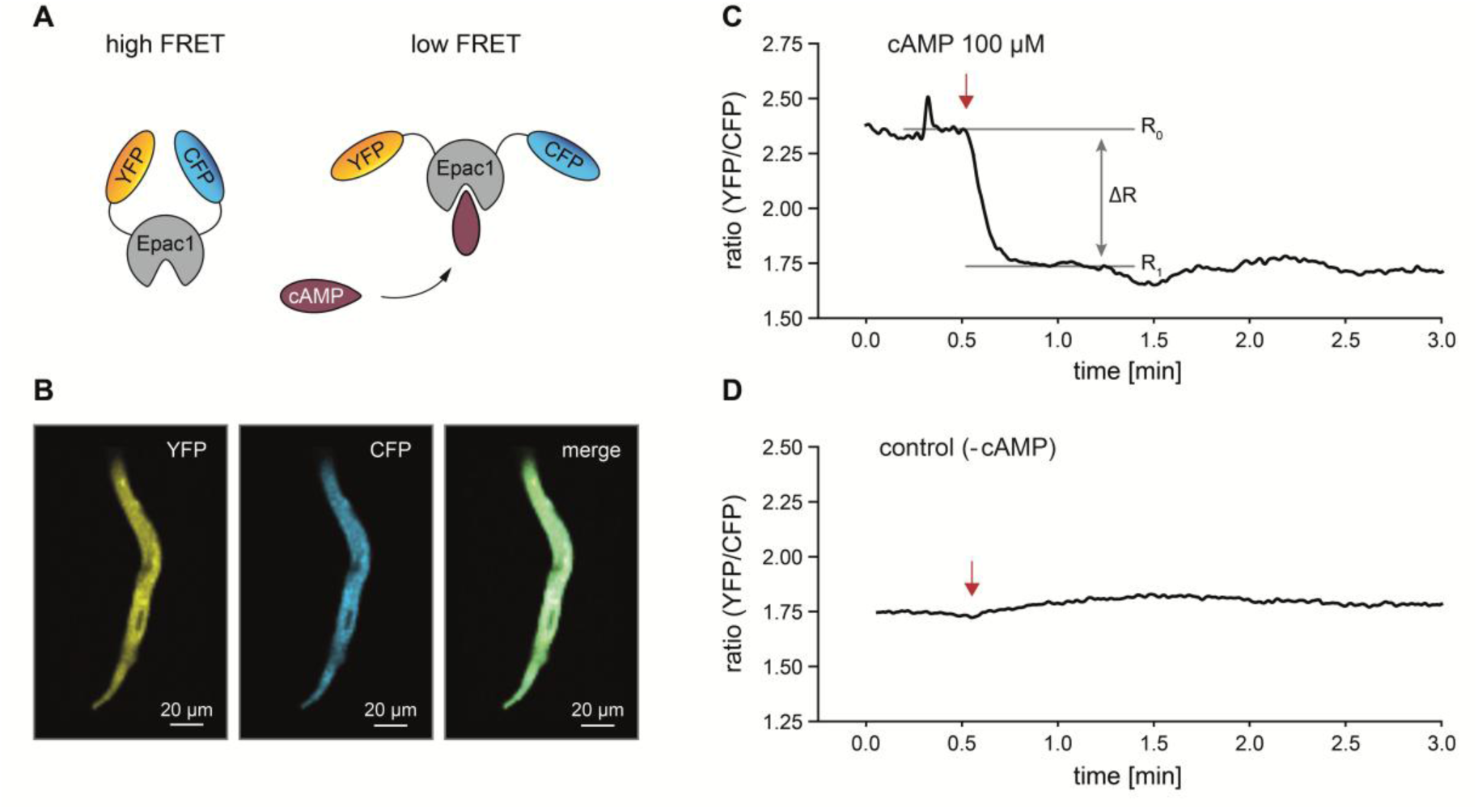
Global and homogenous distribution of cAMP in single SAN cells prepared from mice expressing the Epac1-camps cAMP sensor after cAMP application via the pipette solution in the whole-cell patch clamp configuration. **A** Cartoon of the Epac1-camps cAMP sensor. The cAMP sensor consists of a cAMP binding site from human Epac1 protein (grey), flanked by YFP and CFP. Upon binding of cAMP, a conformational change leads to a spatial separation of CFP and YFP, and a decrease in FRET as indicated by a decrease in YFP (reduction of sensitized emission) and increase in CFP fluorescence (reduction of donor quenching). **B** Fluorescent images of an isolated murine sinoatrial pacemaker cell expressing Epac1-camps. **C, D** The graphs show the time-dependent changes in FRET ratio (YFP/CFP) over time. The total cell area was chosen as the ROI so that changes in the FRET ratio reflect cAMP levels throughout the cytosol. The red arrows indicate the time point of rupture of the membrane patch, i.e., establishment of the whole-cell configuration. A reduction in the FRET ratio is indicative of an increase in intracellular cAMP levels (and vice versa).

Having established effective cAMP application in the whole-cell configuration, we next measured PRCs from SAN cells of WT and HCN4FEA mice (*Figure 5*). Current-clamp recordings of spontaneous action potentials in the absence of cAMP revealed comparable firing rates and MDP in WT and HCN4FEA pacemaker cells (*Figure 5A* and *B, Supplementary Table 2*), demonstrating that baseline characteristics are similar between the two groups of cells. In the presence of cAMP (100 µM), firing rates increased in both groups, and the average firing rates were similar in WT and HCN4FEA cells (*Figure 5E* and *F, Supplementary Table 2*), which is in line with the findings of Fenske et al.^20^ Like in WT, both type 0 and type 1 PRCs were observed in HCN4FEA SAN cells without cAMP, with similar PRC trajectories and proportions compared to WT cells (*Figure 5C* and *D*). Furthermore, in WT and HCN4FEA in the absence of cAMP, the proportion of type 0 PRCs was higher than that of type 1 PRCs (*Figure 5C*). With 100 µM cAMP in the internal recording solution, the proportion of type 1 PRC increased and of type 0 PRCs decreased. Still, the fractions remained similar between genotypes (*Figure 5G*). The shape of type 0 and 1 PRCs was similar in cells from both genotypes in the presence of cAMP (*Figure 5H*). Taken together, these results demonstrate that CDR of HCN4 does not affect phasic entrainment, as firing properties, averaged type 0 and 1 PRCs, and also the proportion of these curves was similar in isolated SAN cells from WT and HCN4FEA mice under both conditions.

**Figure 5.**
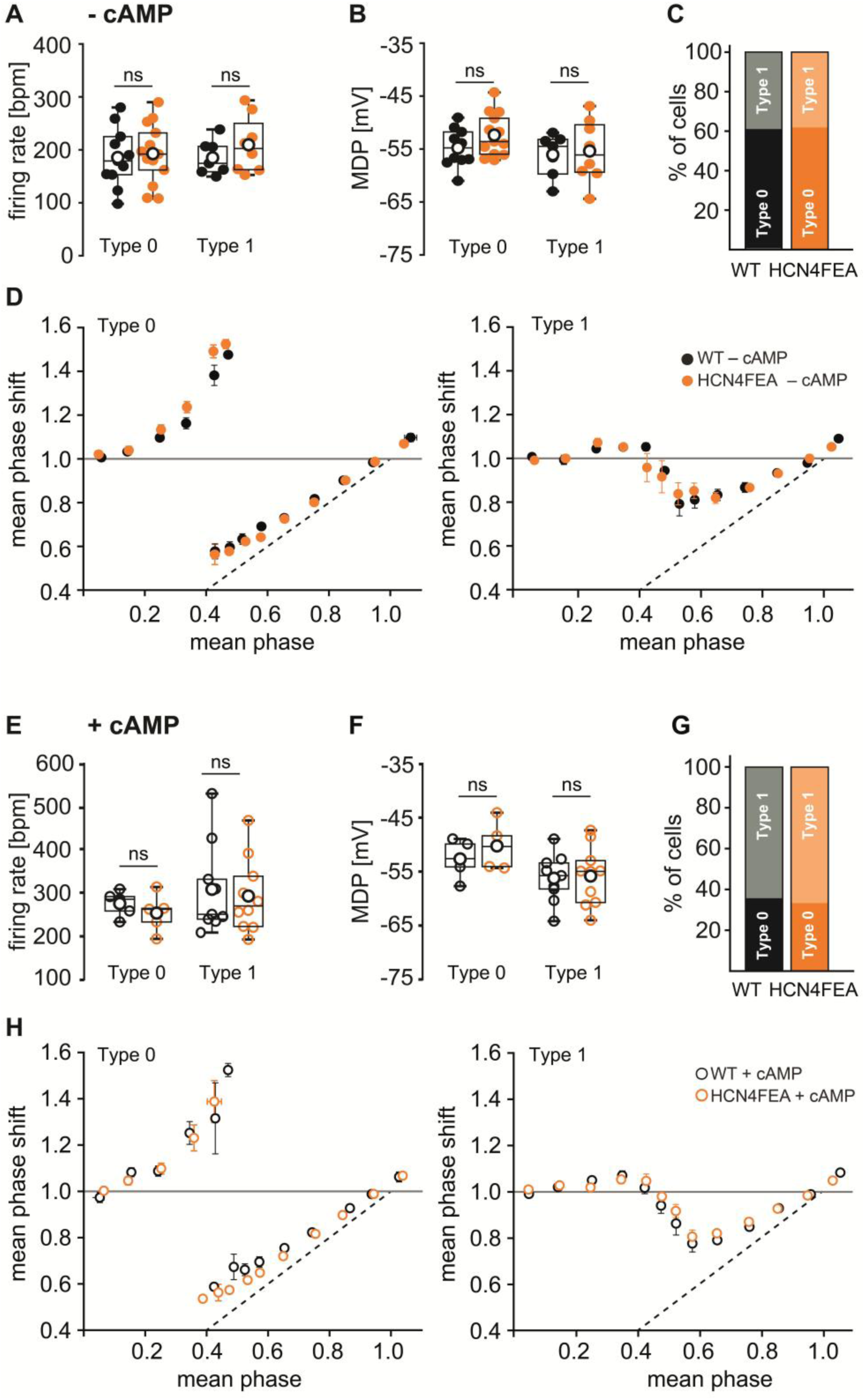
Averaged phase response curves are similar in isolated SAN cells from WT and HCN4FEA mice. *(A-B)* Firing rate and MDP of WT (black closed circles) and HCN4FEA (orange closed circles) cells showing type 0 (left panel) and type 1 (right panel) phase response behaviour during measurements without cAMP in the intracellular solution. *(C)* Percentage of cells showing the typical type 0 or type 1 phase response curves in measurements without cAMP. *(D)* Mean type 0 (left panel) and type 1 (right panel) phase response curves of WT (black closed circles) and HCN4FEA (orange closed circles) SAN cells derived from measurements without cAMP. Dashed lines indicate the line of causality (see text for details). *(E-F)* Firing rate and MDP of WT (black open circles) and HCN4FEA (orange open circles) cells showing type 0 (left panel) and type 1 (right panel) phase response during measurements with cAMP (100 µM) in the intracellular solution. *(G)* Percentage of cells showing the typical type 0 or type 1 phase response curve in measurements in the presence of cAMP. *(H)* Mean type 0 (left panel) and type 1 (right panel) phase response curves of WT (black) and HCN4FEA (orange) SAN cells derived from measurements in the presence of cAMP (100 µM). Dashed lines indicate the line of causality (see text for details). Boxplots show the median line, perc 25/75, and min/max value; open symbols represent the mean value. Significance levels: Holm’s–Sidak post-hoc test following two-way ANOVA (ns, not significant).

### 3.4. Phasic entrainment depends on HCN1 channels in SAN pacemaker cells

Next, PRCs were assessed in SAN cells from HCN1^-/-^ mice and their WT littermates to determine the role of HCN1 channels in phasic entrainment (*Figure 6*). Measurements were made without cAMP, as HCN1 channels are reportedly not significantly regulated by cAMP in SAN cells.^23, 25, 75, 76^ HCN1^-/-^ cells had slower firing rates compared to WT cells (235 ± 10 bpm (n = 28) in HCN1^-/-^ vs. 282 ± 16 bpm (n = 14) in WT; p = 0.012), indicating that HCN1 channels are open during SDD and are crucial for maintaining baseline firing rates, consistent with Fenske et al.^12^ Both type 0 and type 1 PRCs were observed in WT and HCN1^-/-^ SAN cells (*Figure 6B-E*). However, HCN1^-/-^ cells exhibited a higher percentage of type 0 PRCs at 64% compared to 43% in WT cells (*Figure 6E*). Conversely, the proportion of type 1 PRCs in HCN1^-/-^ cells was lower, at 36% compared to 57% in WT cells. The trajectories of type 0 PRCs were similar in WT and HCN1^-/-^ mice (*Figure 6D*, left).

**Figure 6.**
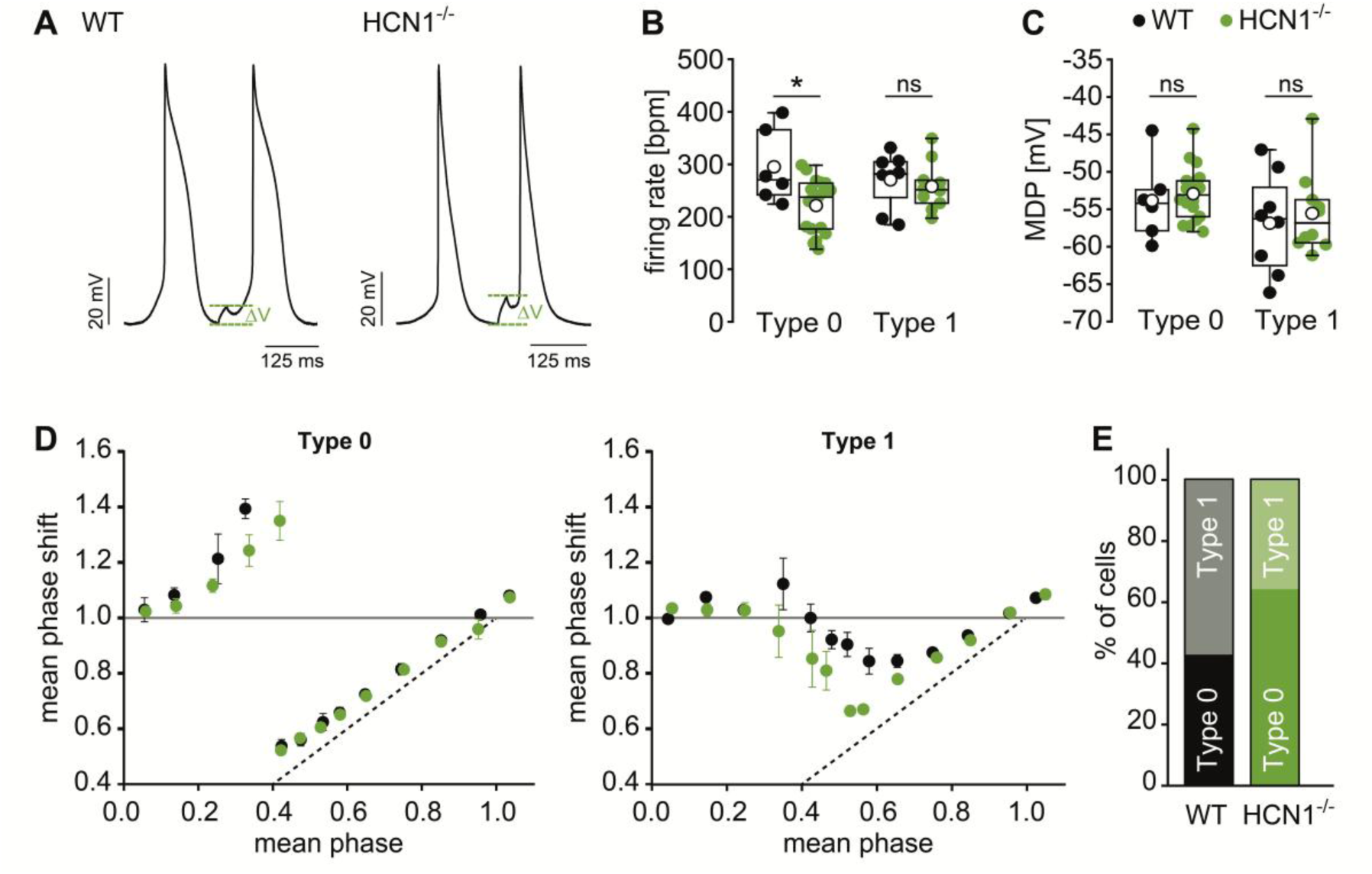
Averaged phase response curves and AP firing rates differ in isolated SAN cells from WT and HCN1^-/-^mice. *(A)* Spontaneous SAN action potential with brief depolarising current pulses (+50 pA, 20 ms). In HCN1^-/-^cells the voltage responses (ΔV) were more pronounced as compared to WT. *(B-C)* Firing rate and MDP of WT (black) and HCN1^-/-^ (green) cells showing type 0 and type 1 phase response behaviour during measurements without cAMP in the intracellular solution. *(D)* Mean type 0 (left panel) and type 1 (right panel) phase response curves of WT (black) and HCN1^-/-^ (green) SAN cells. *(E)* Percentage of all cells showing the typical type 0 or type 1 phase response. Boxplots show the median line, perc 25/75, and min/max value; open symbols represent the mean value. Significance levels: Student’s two-sided t-test (* p < 0.05).

In stark contrast, the shape of type 1 PRCs was different in WT and HCN1^-/-^ cells. Type 1 PRCs of HCN1^-/-^ (versus WT) mice revealed an increased amplitude of phase advance (40.4 ± 5.3 % (n = 10) in HCN1^-/-^ vs. 17.7 ± 2.5 % (n = 8) in WT; p = 0.002), increased area under the curve for phase advances (0.125 ± 0.018 (n = 10) in HCN1^-/-^ vs. 0.048 ± 0.009 (n = 8) in WT; p = 0.003) and increased entrainment range (*Supplementary Table 1*). In addition, the phase advances are shifted towards lower mean stimulation phases along the x-axis, leading to a shift of the unstable equilibrium point from ϕ = 0.41 ± 0.05 (n = 8) in WT cells to ϕ = 0.29 ± 0.03 (n = 10; p = 0.04) in HCN1^-/-^ cells (*Figure 6D*, right; *Supplementary Table 1*). The changes were so pronounced that type 1 PRCs of HCN1^-/-^ cells were nearly monophasic with a marked monophasic peak at a phase of ϕ = 0.53, contrasting with the biphasic PRC in WT cells, that had a more balanced ratio of phase advances and delays. The positions of the stable equilibrium points were similar in WT and HCN1^-/-^ cells. Furthermore, the amplitude of phase delay and the area under the curve for phase delays were unchanged (*Supplementary Table 1*).

The observed differences in type 1 PRCs were qualitatively confirmed in a second, independent data set obtained from perforated-patch experiments (*Supplementary Figure 3*). In these recordings, the mean type 1 PRC of HCN1^-/-^ cells (n = 7) in comparison with the mean type 1 PRC of WT cells (n = 7) had a higher amplitude of phase advance (29.9 % in HCN1^-/-^ vs. 19.7 % in WT), increased area under the curve for phase advances (0.114 in HCN1^-/-^ vs. 0.054 in WT), and a shift of the unstable equilibrium point from ϕ = 0.35 in WT to ϕ = 0.25 in HCN1^-/-^. Like in whole-cell recordings, no differences were observed in the mean type 0 PRCs of WT (n = 11) and HCN1^-/-^ cells (n = 13) measured in the perforated-patch configuration (*Supplementary Figure 3*). These findings rule out the possibility that disturbed intracellular signalling in the whole-cell configuration could interfere with the measurement of PRCs. Interestingly, perforated-patch recordings revealed comparable fractions of type 0 and type 1 PRCs in WT and HCN1^-/-^ SAN cells. Thus, type 0 PRCs do not seem to account for impaired synchronisation in HCN1^-/-^ SANs (see discussion).

To identify the cellular mechanism underlying the increased phase advances in PRCs of HCN1^-/-^ mice as compared to WT mice, the voltage responses (delta V) to premature subthreshold depolarising stimuli of 20 ms were quantified (*Figure 6A*). Given that the difference of the PRC of WT and HCN1^-/-^cells was particularly pronounced in the phase range of 0.25 to 0.5 (*Figure 6D*, right), we calculated delta V for premature stimuli within this range. There was a significant difference in the responses in the phase range between 0.25 and 0.5. In HCN1^-/-^ cells these responses were more pronounced as compared to WT cells, with 14.9 ± 1.7 mV in HCN1^-/-^ (n = 9) and 8.6 ± 1.1 mV in WT cells (n = 8; p = 0.009).

### 3.5. Loss of HCN1 in SAN pacemaker cells leads to reduced synchronisation in the SAN

To explore whether and how the effects of HCN1 and HCN4 on the PRCs translate to SAN network synchronisation and heart rate regularity, we devised a conceptual computational model of the SAN based on the theory of coupled phase oscillators (*Figure 7*, see Methods, Supplementary Material, and ^77^ for details). In brief, unperturbed period durations and PRCs were taken from patch-clamp measurements and imposed on 500 artificial SAN cells, each coupled to 26 nearest neighbours (a cube of length 3). To model the effect of action potential shape, PRCs were convolved with a triangularly shaped model of the murine SAN pacemaker potential, yielding effective PRCs (*Figure 7A*). While effective PRCs hardly differed for type 0 and type 1 cells between WT and HCN4FEA mice, HCN1^-/-^exhibited effective PRCs with monophasic shape and a shift towards lower stimulation phases, indicating a reduced tendency to synchronise. Simulations of the SAN network (*Figure 7B*) showed similar and robust synchronisation behaviour for all conditions except in HCN1^-/-^. The mean rate was slowed down in HCN1^-/-^ mice, and the period was much less regular, as indicated by tachograms and return (Poincaré) maps (*Figure 7C, D*). Thus, a model that only includes information about the phase response behaviour can qualitatively predict experimentally determined synchronisation behaviour^12^ and explains that the reduction of synchronisation ability is due to a leftward shift of type 1 PRCs.

**Figure 7.**
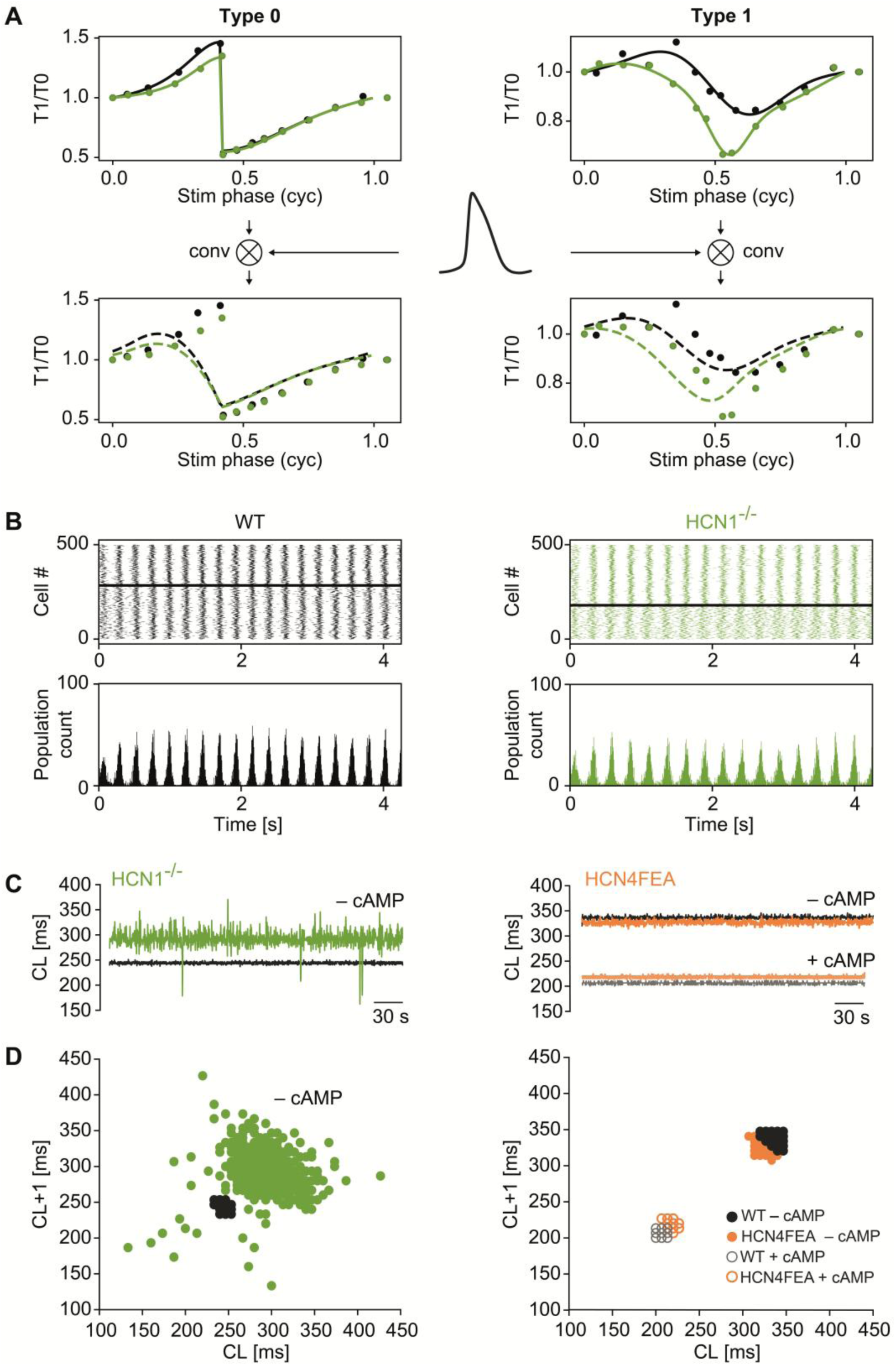
Computer simulations predict increased HR fluctuations in HCN1^-/-^ mice but not in HCN4FEA mice. *(A)* Experimentally determined type 0 and type 1 PRCs (upper panels) were convolved with a murine SAN pacemaker potential model to obtain effective PRCs (lower panels). *(B)* Effective PRCs were used for simulating the synchronisation behaviour of computational models of WT (left) and HCN1^-/-^ (right) SAN networks based on the theory of coupled phase oscillators. These simulations were used to determine the expected cycle lengths over 5 minutes. *(C)* Tachograms were generated by plotting all consecutive cycle lengths (CL) against the time, based on simulations of WT vs. HCN1^-/-^ mice (left) and WT vs. HCN4FEA mice (right). *(D)* Poincaré plots were generated by plotting each cycle length (CL+1) against the previous one (CL) to visualise beat-to-beat dispersion in cycle length. HR fluctuations are more pronounced in HCN1^-/-^ than WT mice. No difference was found between WT and HCN4FEA mice.

Finally, considering that electrical coupling is a key mechanism for SAN synchronization, we performed Calcium imaging experiments on SAN explants of WT mice (*Figure 8* and *Supplementary Videos 2-5*) in control (baseline) and in the presence of gap junction blockers, carbenoxolone (100 µM, *Figure 8A-E, Supplementary Videos 2-3*) or 1-heptanol (10 µM, *Figure 8F-J, Supplementary Videos 4-5*). After perfusion with carbenoxolone or 1-heptanol, the tissues (n=8 for carbenoxolone and n=8 for 1-heptanol) showed a decrease in the frequency of calcium transients (*Figure 8B* and *G*), an increase in cycle length variability (*Figure 8C* and *H*) and an increase in arrhythmic events (*Figure 8D-E* and *I-J*). These results indicate that coupling of single SAN pacemaker cells via gap junctions increases synchrony within the SAN network and stabilizes heart rate.

**Figure 8.**
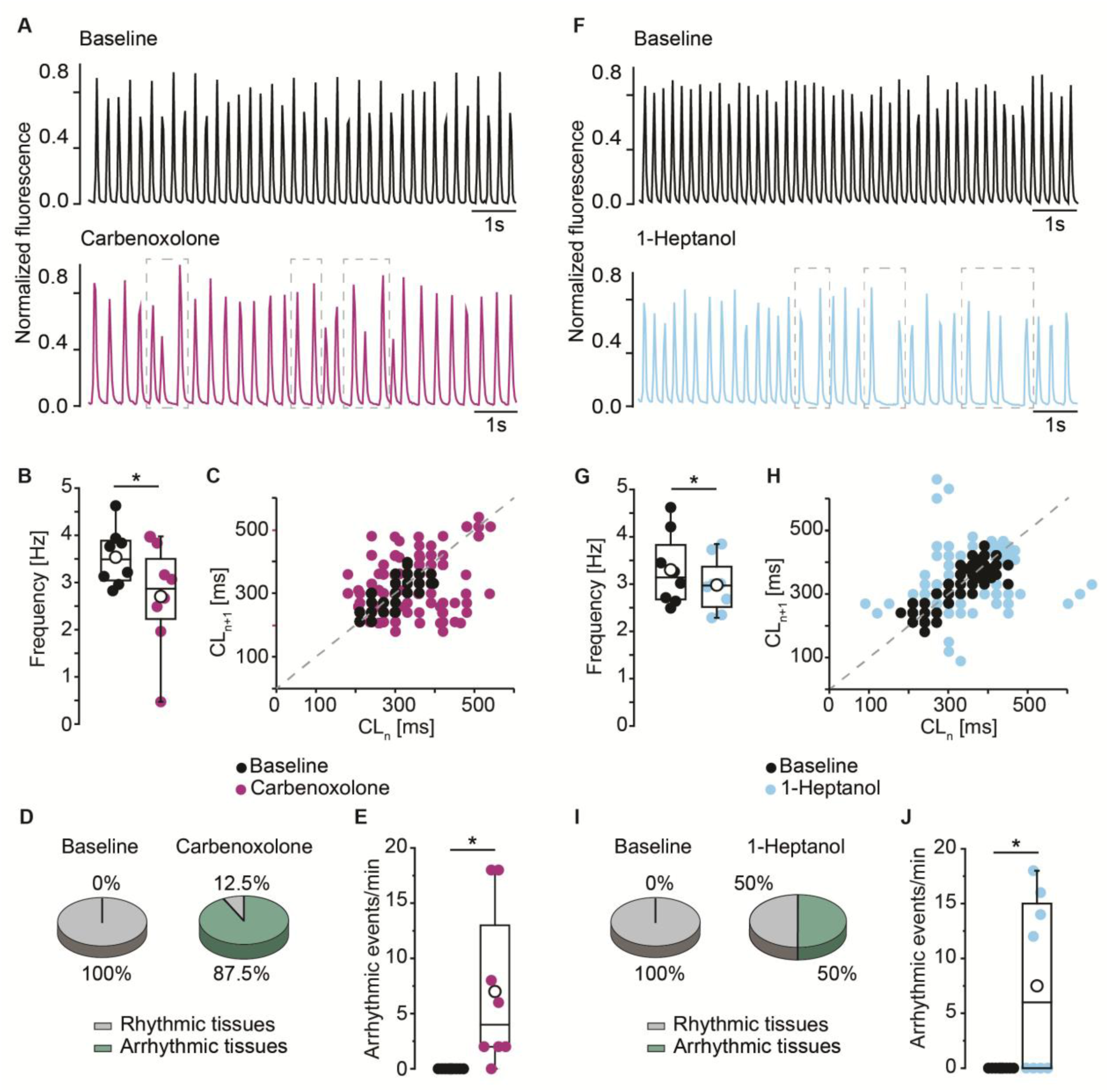
Inhibition of gap junctions induces reduction of frequency, increased variability and arrhythmic events in sinus node explants. (*A*) Representative calcium transients during baseline (black) recording and after perfusion with the gap junction blocker carbenoxolone (100 µM; purple). (*B*) Frequency [Hz] of calcium transients in baseline condition (black circles) and after application of the gap junction blocker carbenoxolone 100 µM (purple circles). (*C*) Poincaré plot of baseline condition (black circles) and after perfusion with Carbenoxolone 100 µM (purple circles) to visualise beat-to-beat dispersion in cycle length (CL). (*D*) Pie charts showing the percentage of tissues presenting arrhythmic events (green) in baseline (left graph) and after perfusion with carbenoxolone (right graph). (*E*) Quantification of arrhythmic events/min in baseline (black) and after inhibition of gap junctions with carbenoxolone (purple). (*F*) Representative calcium transients during baseline (black) recording and after perfusion with the gap junction blocker 1-heptanol (10 µM; light blue). (*G*) Frequency [Hz] of calcium transients in baseline condition (black circles) and after application of the gap junction blocker 1-heptanol 10 µM (light blue circles). (*H*) Poincaré plot of baseline condition (black circles) and after perfusion with 1-heptanol 10 µM (light blue circles. (*I*) Pie charts showing the percentage of tissues presenting arrhythmic events (green) in baseline (left graph) and after perfusion with 1-heptanol (right graph). (*J*) Quantification of arrhythmic events/min in baseline (black) and after inhibition of gap junctions with 1-heptanol (light blue). Boxplots show the median line, perc 25/75, and min/max value; open symbols represent the mean value. Dashed line in the Poincaré plot represents the line of identity. Dashed-line rectangles in the representative calcium transients indicate examples of arrhythmic events. Significance levels: Pair sample t-test.

## 4. Discussion

There are three major advances of the current study. First, PRCs were determined in mouse sinoatrial pacemaker cells. This approach is highly powerful because it allows for predicting the entrainment of the whole SAN network. Second, HCN1 was demonstrated to be crucial for phasic entrainment and synchronisation of the SAN, while CDR of HCN4 is not involved in this process. Third, we propose a mechanism by which HCN1 channels regulate phasic entrainment. These main findings will be discussed in the following.

### 4.1. Characterisation and potential of PRCs in the mouse

To our knowledge, there is no previous study of PRCs in murine SAN pacemaker cells, likely due to the technical challenges posed by the high beating rate of the mouse SAN. Despite these difficulties, PRCs offer several advantages. Most importantly, they encapsulate a pacemaker cell’s entire response to external perturbations. In the native SAN, these external perturbations must be subthreshold but sufficient to alter the pacemaker cycle. These subthreshold signals can arise from pacemaker potentials that are attenuated through high resistance gap junctions between connected pacemaker cells^34^ or from spontaneously occurring low amplitude oscillations in nonfiring pacemaker cells. Another advantage is that PRCs can predict the phasic entrainment of the entire SAN network without the need to experimentally determine numerous individual ionic currents.^59, 60^ Furthermore, the phase-resetting model used in the present study can be fully constrained by up to three free parameters (noise, coupling strength, AP width), making it the most parsimonious model of system behaviour in direct connection to the measured data. It can explain the system behaviour depending on functional properties of SAN cells rather than the underlying subcellular processes that give rise to it, and it is robust against compensatory effects. On the other side, the simulations are based on a simple conceptual model, which represents a region of the SAN, from which cells were isolated, likely including the leading pacemaker region. Anatomical and structural details of the SAN, such as insulation of the SAN periphery from the atria, exit pathways from the SAN, gradients in intercellular electrical coupling by gap junctions, or gradients in ion channel expression are not incorporated in the model.

By utilizing PRCs to predict the network rhythm of the SAN through theoretical modelling, a link can be established between firing at the single cell level and synchronised firing at the level of the SAN network *in vitro* and *in vivo*. This is particularly important in the mouse, as many mouse models for SAN function and SAN dysfunction are available.^6, 12, 16, 18, 20, 21, 78, 79^ Our findings about the role of HCN1 and CDR of HCN4 in the SAN may very likely be also applicable to human SAN physiology and disease mechanisms, as in both, mice and humans, HCN4 and HCN1 are the main isoforms in the SAN, followed by HCN2. However, one needs to keep in mind that HCN1 has a slightly broader distribution in humans than in mice, encompassing all three parts of the human SAN^80, 81^. However, despite shorter APD, different ion channel composition and higher beating rates, PRCs in mice are qualitatively similar to those in other species^34, 40, 82^, making these specific findings relevant for understanding human SAN physiology and disease.

### 4.2. Phasic entrainment involves HCN1, but not CDR of HCN4

Several interesting new characteristics concerning PRCs are determined in pacemaker cells of WT, HCN1^-/-^ and HCN4FEA mice. More type 0 than type 1 PRC were observed for WT SAN cells. The application of cAMP increased the number of cells with type 1 PRCs. This increase is independent of CDR of HCN4 because the proportion of type 0 and type 1 PRCs is similar for WT and HCN4FEA cells with or without cAMP. Importantly, computer simulations predict a regular network rhythm for all conditions (WT and HCN4FEA +/- cAMP). On the other side, there were more type 0 PRCs in SAN pacemaker cells of HCN1^-/-^ mice vs. WT mice. However, this difference was not observed in perforated patch clamp experiments, in which proportions of type 0 and type 1 PRCs were similar between HCN1^-/-^ and WT cells. Still, computer simulations based on perforated-patch data yield an effect comparable to whole-cell conditions (data not shown). Together, this demonstrates that synchronisation is not dependent on the proportions of type 0 and type 1 PRCs, but rather on the qualitative differences in type 1 PRCs of WT and HCN1^-/-^ cells.

Analyses of PRCs revealed similar shapes of WT and HCN4FEA PRCs for both type 0 and type 1 in the absence and presence of cAMP; all parameters determined were comparable (*Supplementary Table 1*). In line with this, computer simulations predicted similar network rhythms for WT and HCN4FEA (*Figure 7C* and *D*). This indicates that the CDR of HCN4 is not involved in the fast, beat-to-beat interactions underlying phasic entrainment. Furthermore, type 0 PRCs recorded in SAN cells isolated from WT and HCN1^-/-^ mice were similar in shape, with all parameters being comparable (*Supplementary Table 1*).

Importantly, the shape of type 1 PRCs differed between WT and HCN1^-/-^ cells. In particular, type 1 PRCs of HCN1^-/-^ cells exhibited a significant leftward shift of phase advances towards lower phases. This shift induces a leftward shift of the unstable repelling fixed point (unstable equilibrium point) at the transition from phase delay to phase advance (*Figure 3*; open purple circle) from ϕ = 0.41 ± 0.05 (n = 8) in WT to ϕ = 0.29 ± 0.03 (n = 10) in HCN1^-/-^ (p = 0.043) (repeller; open purple circle in *Figure 3*).^6, 49, 52, 58, 68, 71–74^ As a result, the range of phases for phase delay is reduced in HCN1^-/-^ and the range available for phase advances is increased. This change is the main factor responsible for the observed increased instability of the network. While there was no significant change in the amplitude and area of phase delay, stimulations at higher phases between ϕ = 0.25 and 0.5 induced greater phase advances in HCN1^-/-^ cells compared to WT cells (maximum of 40.4 ± 5.3 % (n = 10) in HCN1^-/-^ vs. 17.7 ± 2.5 % (n = 8) in WT; p = 0.002). Moreover, the area under the curve was larger for phase advances (0.125 ± 0.018 (n = 10) in HCN1^-/-^ vs. 0.048 ± 0.009 (n = 8) in WT; p = 0.003). Together, these changes result in a monophasic type 1 PRC in HCN1^-/-^ pacemaker cells, characterised by minimal phase delay and a pronounced increase in phase advances with a symmetrical and monophasic maximum centred at ϕ = 0.53.

Altogether, the shift of the unstable equilibrium point, which determines the phase at which the dynamics change from delay to advance, and the monophasic shape of the PRC in HCN1^-/-^ cells reflect an imbalance and loss of flexibility in response to premature impulses. While the increased maximum phase advance indicates a higher propensity for entrainment for longer phases, monophasic PRCs of HCN1^-/-^ can only respond to stimuli by phase advance. As a result, it takes longer for HCN1^-/-^ cells to reach synchrony, and possibly these cells are more prone to respond with inadequately high, i.e. overshooting responses. In contrast, WT cells with biphasic PRC can both advance and delay phases. Finally, the stable equilibrium points at the transition from phase advance to phase delay (*Figure 3*, closed purple circle) are similar in WT and HCN1^-/-^ cells, occurring at a phases of 0 and 1. In conclusion, the reduced ability to synchronise is reflected by the observed changes in type 1 PRCs, while type 0 PRCs do not account for impaired synchronisation in the HCN1^-/-^ SAN.

### 4.3. Mechanism

Our study shows that HCN1, but not CDR of HCN4, is crucial for phasic entrainment. Accordingly, dysfunction of phasic entrainment is present at the single cell level in HCN1^-/-^ mice, contributing to sinus dysrhythmia and severe HR fluctuations observed in a former study in HCN1^-/-^ mice *in vivo.*^12^ In line with this interpretation, HCN1 channels enhance the stability and precision of the heartbeat and reduce beat-to-beat heart rate variability *in vivo.*^12^ The absence of HCN1 impairs these functions. In support of this, computer simulations based on phase response data from HCN1^-/-^ cells predict an irregular network rhythm with high beat-to-beat fluctuations. In contrast, WT conditions predict a regular rhythm (*Figure 7*). Interestingly, blockade of gap junctions (*Figure 8*), key regulators of cell-to-cell synchronization, produces effects similar to those observed in intact SAN preparations from HCN1^-/-^ mice^12^, further supporting the role of HCN1 channels in heart rate stabilization.

To explain the effects of HCN1 on the phase response curve and phasic entrainment,^49^ we suggest the following mechanism: Early in repolarisation, corresponding to a phase range of 0 to 0.25, brief premature subthreshold depolarising stimuli induced similar phase delays in HCN1^-/-^ and WT cells, indicating that HCN1 is not important for inducing phase delays. In contrast, HCN1 plays a key role in phase advances by influencing the early stages of SDD. In line with this, our findings of the present study and also our former studies^12, 83^ suggest that HCN1 channels are open during early SDD. This is due to the characteristic deactivation kinetics of HCN1 which is very slow in relation to the duration of an action potential (which is rather short) and, hence, too slow to close HCN1 channels during the time course of a pacemaker potential. During late repolarisation, the membrane potential gradually falls below −40 mV (the reversal potential of HCN channels) and increases the driving force for a depolarising inward current through open HCN1 channels. In WT, open HCN1 channels decrease the total membrane resistance. The decrease in membrane resistance due to open HCN1 makes it more difficult for external stimuli to induce depolarisations and a phase advance. Reversely, the absence of HCN1 would increase membrane resistance, leading to larger voltage amplitudes in response to depolarising stimuli. Indeed, our results demonstrate that in HCN1^-/-^ SAN pacemaker cells, depolarising subthreshold current stimuli (20 ms, phase ranging between 0.25 and 0.5) early in the SDD induce a larger voltage response (delta Vm) than in their WT counterparts (*Figure 6A*).

Interestingly, in a previous study, we show that open HCN1 channels at the resting membrane potential result in more positive MDP in WT (vs HCN1^-/-^) cells.^12^ While there is a trend toward more positive MDP in WT as compared to HCN1^-/-^ cells, this effect is not significant in the cell population investigated in the present study. Possible reasons could be that the former study focused on elongated cells, while in the present study, short spindle and elongated SAN cell types were investigated. In any case, a more positive MDP in WT cells (vs HCN1^-/-^ cells) would facilitate the effects of reduced membrane resistance. The more positive MDP would inactivate a fraction of Na^+^ and Ca^2+^ channels. As a result, during the following SDD, fewer Na^+^ and Ca^2+^ channels would be available for inducing a phase advance in response to a premature stimulation. Reversely, the absence of HCN1 would result in a more negative MDP, leading to more available Na^+^ and Ca^2+^ channels and as a consequence a faster response to premature stimulation late in SDD and a greater phase advance.

Taken together, we provide new evidence highlighting the distinct roles of the two primary HCN channel isoforms in the SAN. Both HCN1 and HCN4 are crucial for intrinsic entrainment, stabilising the SAN network, generating a rhythmic heartbeat, and reducing beat-to-beat heart rate variability (*Figure 1*).^12, 20^ However, their contributions differ (*Figure 1*). cAMP-dependent regulation of HCN4 is involved in tonic entrainment between firing and nonfiring pacemaker cells, maintaining a balance between inhibition by nonfiring cells and excitation by firing cells in the SAN through long-lasting interactions.^6,20^ In contrast, HCN1 is involved in phasic entrainment, synchronising the firing rates of adjacent pacemaker cells in the firing mode on a beat-to-beat basis through brief interactions in the diastolic voltage range. In the HCN1^-/-^ mouse, physiological stimuli could trigger an erroneous and overshooting response via gap junctions by inadequate phase shortening or phase lengthening. At the network level, this would lead to a defective entrainment process, divergence of the frequencies of individual pacemaker cells, and an unstable heart rate with large fluctuations.

This work enhances our understanding of the pacemaker process at the single-cell and tissue levels and the mechanisms underlying SAN function and dysfunction *in vivo*. It thereby opens the door for several new clinically relevant applications. Understanding how I_f_ blockers may affect heart rate and rhythm stability is crucial for assessing potential side effects of current and future subtype-specific HCN channel inhibitors. Moreover, our findings could support new diagnostic strategies for detecting patients at risk of SAN dysfunction. The knowledge that I_f_ inhibition could potentially lead to sinus node dysrhythmia could be used for assessment of the “diastolic depolarization reserve” by pharmacological provocation tests with selective HCN channel blockers (or ivabradine), analogous to the Sotalol test (assessment of the repolarization reserve for long QT syndrome^84^). Another diagnostic test for SND could be genetic sequencing of HCN channel variants.

Ultimately, these findings pave the way for innovative therapeutic approaches, including targeted channel modulation and future cell or gene therapies to restore pacemaker stability.

## Authors’ contributions

Conceptualization: CWS, SF, MB, CL; Methodology: CWS, SF, KH, CL, VON; Investigation: KH, CP, MO, LP, MS, NA, MH, CL, YW, RR, CF, MGP, DK; Formal analysis: KH, CP, MO, LP, MS, NA, MH, CF, CL, SF, CWS; Writing, original draft: CWS, KH, SF, CL; Writing, review and editing: CWS, KH, SF, CL, CP, LP, NA, AK, JR, BA; Visualization: SF, KH, CP, NA, RR; Project administration: CWS, SF; Funding Acquisition; CWS, SF, MB

## Funding acknowledgments

This work was supported by the German Research Foundation (DFG): WA 2597/3-2; TRR 152, P28; FE 1929/2-2; FE 1929/1-1; BI484/5-2; INST 192/543-1 FUGG, CWS

## Data availability

The data underlying this article will be shared on reasonable request to the corresponding authors.

## References

1. Billman GE. Heart rate variability - a historical perspective. Front Physiol. 2011;2:86

2. Mesirca P, Fedorov VV, Hund TJ, Torrente AG, Bidaud I, Mohler PJ, Mangoni ME. Pharmacologic approach to sinoatrial node dysfunction. Annu Rev Pharmacol Toxicol. 2021;61:757–778

3. De Ponti R, Marazzato J, Bagliani G, Leonelli FM, Padeletti L. Sick sinus syndrome. Card Electrophysiol Clin. 2018;10:183–195

4. Dobrzynski H, Boyett MR, Anderson RH. New insights into pacemaker activity: Promoting understanding of sick sinus syndrome. Circulation. 2007;115:1921–1932

5. Wallace MJ, El Refaey M, Mesirca P, Hund TJ, Mangoni ME, Mohler PJ. Genetic complexity of sinoatrial node dysfunction. Front Genet. 2021;12:654925

6. Hennis K, Piantoni C, Biel M, Fenske S, Wahl-Schott C. Pacemaker channels and the chronotropic response in health and disease. Circ Res. 2024;134:1348–1378

7. Lakatta EG, Maltsev VA, Vinogradova TM. A coupled system of intracellular ca2+ clocks and surface membrane voltage clocks controls the timekeeping mechanism of the heart’s pacemaker. Circ Res. 2010;106:659–673

8. Hennis K, Rotzer RD, Piantoni C, Biel M, Wahl-Schott C, Fenske S. Speeding up the heart? Traditional and new perspectives on hcn4 function. Front Physiol. 2021;12:669029

9. Mangoni ME, Nargeot J. Genesis and regulation of the heart automaticity. Physiological reviews. 2008;88:919–982

10. DiFrancesco D. Pacemaker mechanisms in cardiac tissue. Annu Rev Physiol. 1993;55:455–472

11. Brown HF, DiFrancesco D, Noble SJ. How does adrenaline accelerate the heart? Nature. 1979;280:235–236

12. Fenske S, Krause SC, Hassan SI, Becirovic E, Auer F, Bernard R, Kupatt C, Lange P, Ziegler T, Wotjak CT, Zhang H, Hammelmann V, Paparizos C, Biel M, Wahl-Schott CA. Sick sinus syndrome in hcn1-deficient mice. Circulation. 2013;128:2585–2594

13. Hennis K, Biel M, Fenske S, Wahl-Schott C. Paradigm shift: New concepts for hcn4 function in cardiac pacemaking. Pflugers Arch. 2022;474:649–663

14. Marionneau C, Couette B, Liu J, Li H, Mangoni ME, Nargeot J, Lei M, Escande D, Demolombe S. Specific pattern of ionic channel gene expression associated with pacemaker activity in the mouse heart. J Physiol. 2005;562:223–234

15. Herrmann S, Layh B, Ludwig A. Novel insights into the distribution of cardiac hcn channels: An expression study in the mouse heart. J Mol Cell Cardiol. 2011;51:997–1006

16. Baruscotti M, Bucchi A, Viscomi C, Mandelli G, Consalez G, Gnecchi-Rusconi T, Montano N, Casali KR, Micheloni S, Barbuti A, DiFrancesco D. Deep bradycardia and heart block caused by inducible cardiac-specific knockout of the pacemaker channel gene hcn4. Proceedings of the National Academy of Sciences of the United States of America. 2011;108:1705–1710

17. Stieber J, Herrmann S, Feil S, Loster J, Feil R, Biel M, Hofmann F, Ludwig A. The hyperpolarization-activated channel hcn4 is required for the generation of pacemaker action potentials in the embryonic heart. Proceedings of the National Academy of Sciences of the United States of America. 2003;100:15235–15240

18. Herrmann S, Stieber J, Stockl G, Hofmann F, Ludwig A. Hcn4 provides a ‘depolarization reserve’ and is not required for heart rate acceleration in mice. EMBO J. 2007;26:4423–4432

19. Hoesl E, Stieber J, Herrmann S, Feil S, Tybl E, Hofmann F, Feil R, Ludwig A. Tamoxifen-inducible gene deletion in the cardiac conduction system. J Mol Cell Cardiol. 2008;45:62–69

20. Fenske S, Hennis K, Rotzer RD, Brox VF, Becirovic E, Scharr A, Gruner C, Ziegler T, Mehlfeld V, Brennan J, Efimov IR, Pauza AG, Moser M, Wotjak CT, Kupatt C, Gonner R, Zhang R, Zhang H, Zong X, Biel M, Wahl-Schott C. Camp-dependent regulation of hcn4 controls the tonic entrainment process in sinoatrial node pacemaker cells. Nat Commun. 2020;11:5555

21. Mesirca P, Alig J, Torrente AG, Muller JC, Marger L, Rollin A, Marquilly C, Vincent A, Dubel S, Bidaud I, Fernandez A, Seniuk A, Engeland B, Singh J, Miquerol L, Ehmke H, Eschenhagen T, Nargeot J, Wickman K, Isbrandt D, Mangoni ME. Cardiac arrhythmia induced by genetic silencing of ‘funny’ (f) channels is rescued by girk4 inactivation. Nat Commun. 2014;5:4664

22. DiFrancesco D, Tortora P. Direct activation of cardiac pacemaker channels by intracellular cyclic amp. Nature. 1991;351:145–147

23. Wainger BJ, DeGennaro M, Santoro B, Siegelbaum SA, Tibbs GR. Molecular mechanism of camp modulation of hcn pacemaker channels. Nature. 2001;411:805–810

24. Biel M, Wahl-Schott C, Michalakis S, Zong X. Hyperpolarization-activated cation channels: From genes to function. Physiological reviews. 2009;89:847–885

25. Biel M, Schneider A, Wahl C. Cardiac hcn channels: Structure, function, and modulation. Trends Cardiovasc Med. 2002;12:206–212

26. Xu X, Vysotskaya ZV, Liu Q, Zhou L. Structural basis for the camp-dependent gating in the human hcn4 channel. The Journal of biological chemistry. 2010;285:37082–37091

27. Saponaro A, Cantini F, Porro A, Bucchi A, DiFrancesco D, Maione V, Donadoni C, Introini B, Mesirca P, Mangoni ME, Thiel G, Banci L, Santoro B, Moroni A. A synthetic peptide that prevents camp regulation in mammalian hyperpolarization-activated cyclic nucleotide-gated (hcn) channels. eLife. 2018;7

28. Boyett MR, Honjo H, Kodama I. The sinoatrial node, a heterogeneous pacemaker structure. Cardiovasc Res. 2000;47:658–687

29. Fedorov VV, Glukhov AV, Chang R, Kostecki G, Aferol H, Hucker WJ, Wuskell JP, Loew LM, Schuessler RB, Moazami N, Efimov IR. Optical mapping of the isolated coronary-perfused human sinus node. J Am Coll Cardiol. 2010;56:1386–1394

30. Boineau JP, Schuessler RB, Roeske WR, Autry LJ, Miller CB, Wylds AC. Quantitative relation between sites of atrial impulse origin and cycle length. Am J Physiol. 1983;245:H781–789

31. Shibata N, Inada S, Mitsui K, Honjo H, Yamamoto M, Niwa R, Boyett MR, Kodama I. Pacemaker shift in the rabbit sinoatrial node in response to vagal nerve stimulation. Exp Physiol. 2001;86:177–184

32. Glukhov AV, Fedorov VV, Anderson ME, Mohler PJ, Efimov IR. Functional anatomy of the murine sinus node: High-resolution optical mapping of ankyrin-b heterozygous mice. Am J Physiol Heart Circ Physiol. 2010;299:H482–491

33. Schuessler RB, Boineau JP, Bromberg BI. Origin of the sinus impulse. J Cardiovasc Electrophysiol. 1996;7:263–274

34. Jalife J. Mutual entrainment and electrical coupling as mechanisms for synchronous firing of rabbit sino-atrial pace-maker cells. J Physiol. 1984;356:221–243

35. Clancy CE, Santana LF. Evolving discovery of the origin of the heartbeat: A new perspective on sinus rhythm. JACC Clin Electrophysiol. 2020;6:932–934

36. Grainger N, Guarina L, Cudmore RH, Santana LF. The organization of the sinoatrial node microvasculature varies regionally to match local myocyte excitability. Function (Oxf). 2021;2:zqab031

37. Maltsev VA, Stern MD. The paradigm shift: Heartbeat initiation without “the pacemaker cell”. Front Physiol. 2022;13:1090162

38. Bychkov R, Juhaszova M, Tsutsui K, Coletta C, Stern MD, Maltsev VA, Lakatta EG. Synchronized cardiac impulses emerge from heterogeneous local calcium signals within and among cells of pacemaker tissue. JACC Clin Electrophysiol. 2020;6:907–931

39. Hennis K, Biel M, Wahl-Schott C, Fenske S. Beyond pacemaking: Hcn channels in sinoatrial node function. Prog Biophys Mol Biol. 2021;166:51–60

40. Jalife J, Moe GK. Phasic effects of vagal stimulation on pacemaker activity of the isolated sinus node of the young cat. Circ Res. 1979;45:595–608

41. Michaels DC, Matyas EP, Jalife J. Mechanisms of sinoatrial pacemaker synchronization: A new hypothesis. Circ Res. 1987;61:704–714

42. DiFrancesco D, Ferroni A, Mazzanti M, Tromba C. Properties of the hyperpolarizing-activated current (if) in cells isolated from the rabbit sino-atrial node. J Physiol. 1986;377:61–88

43. Magee JC. Dendritic hyperpolarization-activated currents modify the integrative properties of hippocampal ca1 pyramidal neurons. J Neurosci. 1998;18:7613–7624

44. Mishra P, Narayanan R. The enigmatic hcn channels: A cellular neurophysiology perspective. Proteins. 2025;93:72–92

45. Narayanan R, Johnston D. The h channel mediates location dependence and plasticity of intrinsic phase response in rat hippocampal neurons. J Neurosci. 2008;28:5846–5860

46. Nolan MF, Dudman JT, Dodson PD, Santoro B. Hcn1 channels control resting and active integrative properties of stellate cells from layer ii of the entorhinal cortex. J Neurosci. 2007;27:12440–12451

47. Nolan MF, Malleret G, Dudman JT, Buhl DL, Santoro B, Gibbs E, Vronskaya S, Buzsaki G, Siegelbaum SA, Kandel ER, Morozov A. A behavioral role for dendritic integration: Hcn1 channels constrain spatial memory and plasticity at inputs to distal dendrites of ca1 pyramidal neurons. Cell. 2004;119:719–732

48. Robinson RB, Siegelbaum SA. Hyperpolarization-activated cation currents: From molecules to physiological function. Annu Rev Physiol. 2003;65:453–480

49. Michaels DC, Matyas EP, Jalife J. Dynamic interactions and mutual synchronization of sinoatrial node pacemaker cells. A mathematical model. Circ Res. 1986;58:706–720

50. Calebiro D, Nikolaev VO, Gagliani MC, de Filippis T, Dees C, Tacchetti C, Persani L, Lohse MJ. Persistent camp-signals triggered by internalized g-protein-coupled receptors. PLoS Biol. 2009;7:e1000172

51. Fenske S, Probstle R, Auer F, Hassan S, Marks V, Pauza DH, Biel M, Wahl-Schott C. Comprehensive multilevel in vivo and in vitro analysis of heart rate fluctuations in mice by ecg telemetry and electrophysiology. Nat Protoc. 2016;11:61–86

52. Granada A, Hennig RM, Ronacher B, Kramer A, Herzel H. Phase response curves elucidating the dynamics of coupled oscillators. Methods Enzymol. 2009;454:1–27

53. Jimenez A, Lu Y, Jambhekar A, Lahav G. Principles, mechanisms and functions of entrainment in biological oscillators. Interface Focus. 2022;12:20210088

54. Netoff TI. Phase response curve, measurement, and shape of general. Encyclopedia of Computational Neuroscience. 2014

55. Tsalikakis DG, Zhang HG, Fotiadis DI, Kremmydas GP, Michalis LK. Phase response characteristics of sinoatrial node cells. Comput Biol Med. 2007;37:8–20

56. Van Meerwijk WP, deBruin G, Van Ginneken CG, VanHartevelt J, Jongsma HJ, Kruyt EW, Scott SS, Ypey DL. Phase resetting properties of cardiac pacemaker cells. J Gen Physiol. 1984;83:613–629

57. Coster AC, Celler BG. Phase response of model sinoatrial node cells. Ann Biomed Eng. 2003;31:271–283

58. Ikeda N. Model of bidirectional interaction between myocardial pacemakers based on the phase response curve. Biol Cybern. 1982;43:157–167

59. Ermentrout GB, Beverlin B, Netoff T. Phase response curves to measure ion channel effects on neurons. In: Schultheiss NW, Prinz AA, Butera RJ, eds. Phase response curves in neuroscience: Theory, experiment, and analysis. New York, NY: Springer New York; 2012:207-236.

60. Granada AE, Bordyugov G, Kramer A, Herzel H. Human chronotypes from a theoretical perspective. PLoS One. 2013;8:e59464

61. Hennis K, Rotzer RD, Rilling J, Wu Y, Thalhammer SB, Biel M, Wahl-Schott C, Fenske S. In vivo and ex vivo electrophysiological study of the mouse heart to characterize the cardiac conduction system, including atrial and ventricular vulnerability. Nat Protoc. 2022;17:1189–1222

62. Brennan JA, Chen Q, Gams A, Dyavanapalli J, Mendelowitz D, Peng W, Efimov IR. Evidence of superior and inferior sinoatrial nodes in the mammalian heart. JACC Clin Electrophysiol. 2020;6:1827–1840

63. James TN. Anatomy of the cardiac conduction system in the rabbit. Circ Res. 1967;20:638–648

64. Kagan CM, Amara RS, Haq M, Dickfeld TM, See VY, Shorofsky SR. Bachmann’s bundle’s unique physiology: Reviewing how it made an atypical flutter even more atypical. JACC Case Rep. 2023;9:101591

65. van Campenhout MJ, Yaksh A, Kik C, de Jaegere PP, Ho SY, Allessie MA, de Groot NM. Bachmann’s bundle: A key player in the development of atrial fibrillation? Circ Arrhythm Electrophysiol. 2013;6:1041–1046

66. Infeld M, Lobel R, Hopper M, Habel N, Winget J, Correa de Sa D, Thompson N, Sanchez-Quintana D, Lustgarten D. Biatrial resynchronization with electrogram-guided bachmann bundle pacing. JACC Clin Electrophysiol. 2024;10:2103–2107

67. Cavero I, Holzgrefe H. Internodal conduction pathways: Revisiting a century-long debate on their existence, morphology, and location in the context of 2023 best science. Adv Physiol Educ. 2023;47:838–850

68. Ikeda N, Tsuruta H, Sato T. Difference equation model of the entrainment of myocardial pacemaker cells based on the phase response curve. Biol Cybern. 1981;42:117–128

69. Oprisan SA, Boutan C. Prediction of entrainment and 1:1 phase-locked modes in two-neuron networks based on the phase resetting curve method. Int J Neurosci. 2008;118:867–890

70. Anumonwo JM, Delmar M, Vinet A, Michaels DC, Jalife J. Phase resetting and entrainment of pacemaker activity in single sinus nodal cells. Circ Res. 1991;68:1138–1153

71. Pfeuty B, Thommen Q, Lefranc M. Robust entrainment of circadian oscillators requires specific phase response curves. Biophys J. 2011;100:2557–2565

72. Ikeda N, Yamamoto H, Sato T. Pathology of the pacemaker network. Mathematical Modelling. 1986;7:889–904

73. Dexter F, Levy MN, Rudy Y. Mathematical model of the changes in heart rate elicited by vagal stimulation. Circ Res. 1989;65:1330–1339

74. May RM. Simple mathematical models with very complicated dynamics. Nature. 1976;261:459–467

75. Chen S, Wang J, Siegelbaum SA. Properties of hyperpolarization-activated pacemaker current defined by coassembly of hcn1 and hcn2 subunits and basal modulation by cyclic nucleotide. J Gen Physiol. 2001;117:491–504

76. Moroni A, Gorza L, Beltrame M, Gravante B, Vaccari T, Bianchi ME, Altomare C, Longhi R, Heurteaux C, Vitadello M, Malgaroli A, DiFrancesco D. Hyperpolarization-activated cyclic nucleotide-gated channel 1 is a molecular determinant of the cardiac pacemaker current i(f). The Journal of biological chemistry. 2001;276:29233–29241

77. Winfree AT. Biological rhythms and the behavior of populations of coupled oscillators. J Theor Biol. 1967;16:15–42

78. Kozasa Y, Nakashima N, Ito M, Ishikawa T, Kimoto H, Ushijima K, Makita N, Takano M. Hcn4 pacemaker channels attenuate the parasympathetic response and stabilize the spontaneous firing of the sinoatrial node. J Physiol. 2018;596:809–825

79. Alig J, Marger L, Mesirca P, Ehmke H, Mangoni ME, Isbrandt D. Control of heart rate by camp sensitivity of hcn channels. Proceedings of the National Academy of Sciences of the United States of America. 2009;106:12189–12194

80. Li N, Artiga E, Kalyanasundaram A, Hansen BJ, Webb A, Pietrzak M, Biesiadecki B, Whitson B, Mokadam NA, Janssen PML, Hummel JD, Mohler PJ, Dobrzynski H, Fedorov VV. Altered microrna and mrna profiles during heart failure in the human sinoatrial node. Sci Rep. 2021;11:19328

81. Li N, Csepe TA, Hansen BJ, Dobrzynski H, Higgins RS, Kilic A, Mohler PJ, Janssen PM, Rosen MR, Biesiadecki BJ, Fedorov VV. Molecular mapping of sinoatrial node hcn channel expression in the human heart. Circ Arrhythm Electrophysiol. 2015;8:1219–1227

82. Verheijck EE, Wilders R, Joyner RW, Golod DA, Kumar R, Jongsma HJ, Bouman LN, van Ginneken AC. Pacemaker synchronization of electrically coupled rabbit sinoatrial node cells. J Gen Physiol. 1998;111:95–112

83. Fenske S, Krause S, Biel M, Wahl-Schott C. The role of hcn channels in ventricular repolarization. Trends Cardiovasc Med. 2011;21:216–220

84. Kaab S, Hinterseer M, Nabauer M, Steinbeck G. Sotalol testing unmasks altered repolarization in patients with suspected acquired long-qt-syndrome--a case-control pilot study using i.V. Sotalol. Eur Heart J. 2003;24:649–657

