## Supplementary Information for "Differential contribution of HCN1 and HCN4 to the synchronisation of sinoatrial pacemaker cells"

* Shared first authorship
[#] Shared last authorship

**Short title: Synchronisation in SAN cells**

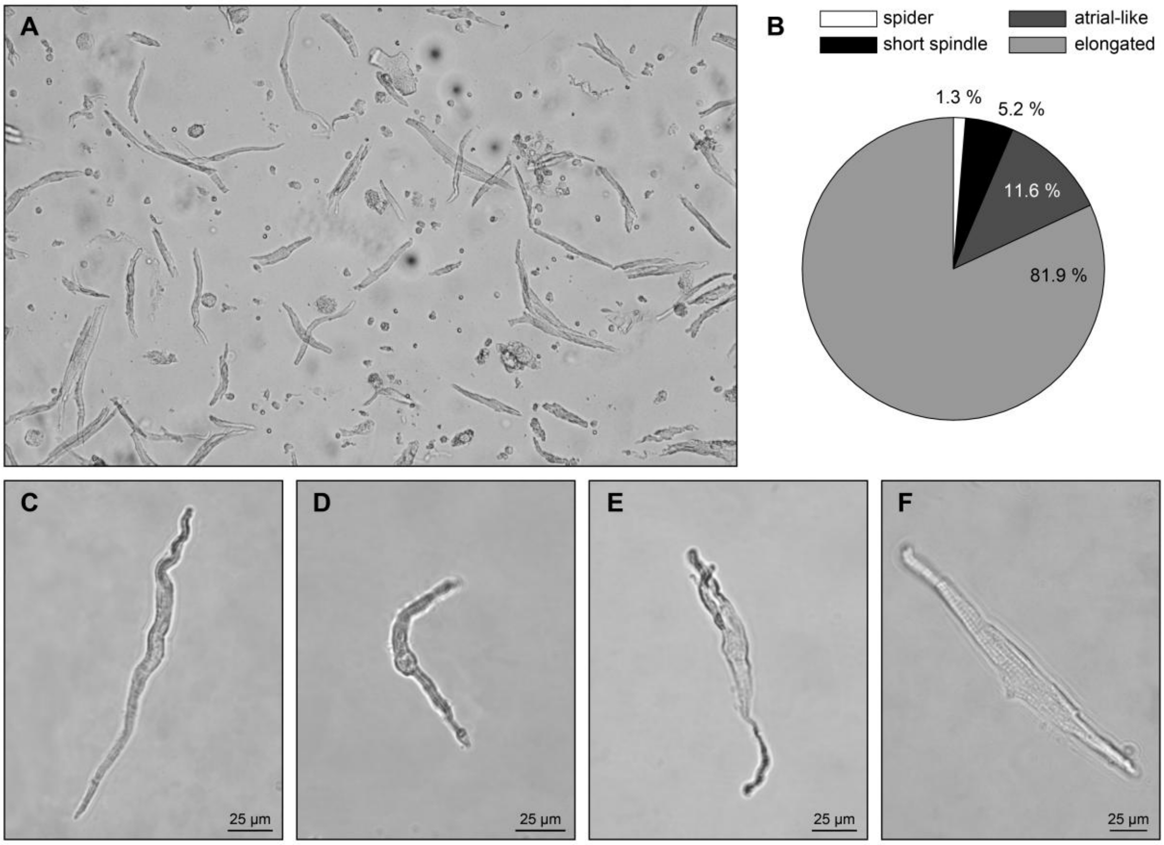


**Supplementary Figure 1** Single-cell isolations from the SAN yield different subtypes of pacemaker cells. *(A)* Representative image of isolated SAN cells attached to a PLL-coated coverslip and placed in the perfusion chamber of the electrophysiological recording setup. *(B)* Quantification of the different pacemaker cell subtypes (as shown in C-F) after readaptation to a physiological extracellular $Ca^{2+}$ concentration (1.8 mM). A total of 2888 $Ca^{2+}$-tolerant cells from 29 mice were visually counted and categorized into subgroups based on cell morphology. *(C-F)* Representative images of an elongated spindle cell (C), short spindle cell (D), spider cell (E), and atrial-like cell (F). Atrial-like cells and spider cells were identified by visual inspection and not included in the analysis.

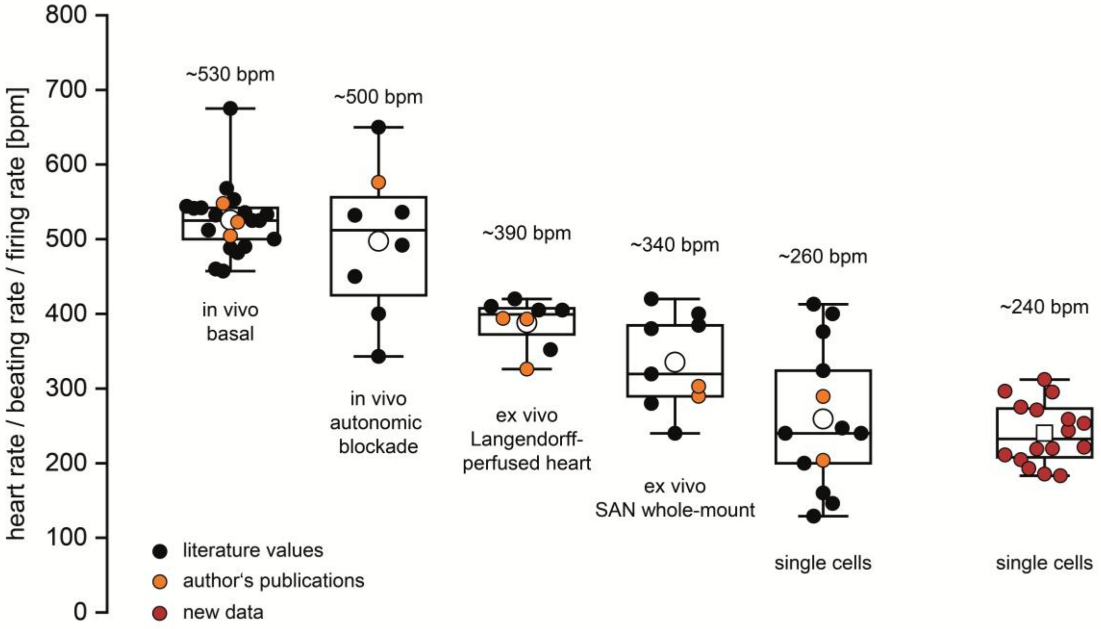


**Supplementary Figure 2** Screening of published data from mice (29 publications from 1998 – 2025) reveals gradually decreasing beating rates from the in vivo level down to the single cell level[1-29]. Previously published data by the authors of the present study are highlighted in orange. New data included in the present study are highlighted in red. Boxplots show the median line, perc 25/75, and min/max value; open symbols represent the mean value.

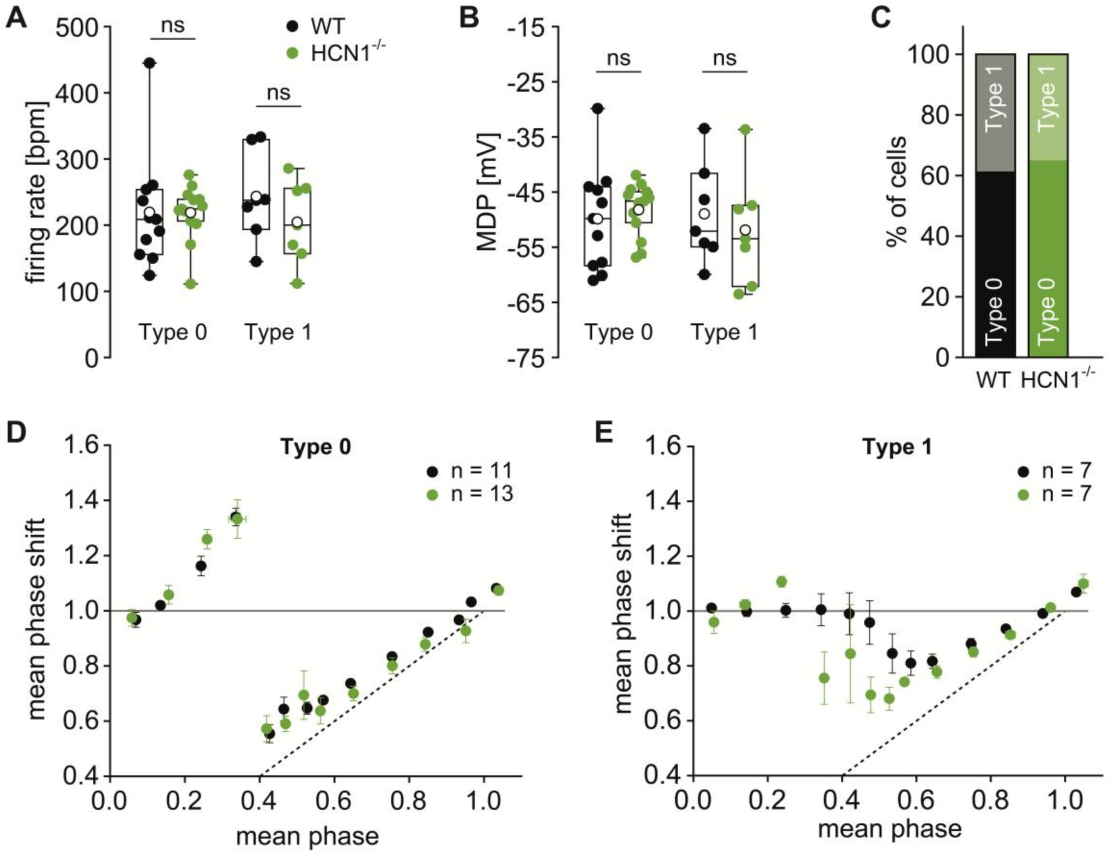


**Supplementary Figure 3** Perforated patch-clamp recordings with amphotericin B qualitatively confirm the differences in PRCs of WT and HCN1$^{-/-}$ SAN cells measured in the whole-cell configuration (see main manuscript). *(A-B)* Firing rate and MDP of WT (black) and HCN1$^{-/-}$ (green) cells showing type 0 and type 1 phase response behaviour during measurements in the perforated-patch configuration. *(C)* Percentage of all cells showing the typical type 0 or type 1 phase response. *(D)* Mean type 0 phase response curves of WT (black) and HCN1$^{-/-}$ (green) SAN cells. *(E)* Mean type 1 phase response curves of WT (black) and HCN1$^{-/-}$ (green) SAN cells. Boxplots show the median line, perc 25/75, and min/max value; open symbols represent the mean value. Significance levels: Student's two-sided t-test (ns: $p > 0.05$).

### Description of additional supplementary material

**Supplementary Table 1** Separate Excel table containing PRC parameters of the experimental data measured in the whole-cell configuration as shown in Figures 4-5 in the main manuscript. To derive PRC parameters, model functions were fitted to the experimental data of each cell as described in the Model Supplement.

**Supplementary Table 2** Separate Excel table containing action potential (AP) parameters of all SAN cells investigated in this study. These data comprise the firing rate, peak voltage (Vmax), maximum diastolic potential (MDP), slope of slow diastolic depolarization (SDD), and action potential duration (APD, measured from the peak to 10%, 25%, 50%, 75%, 90% of repolarisation). APs of mutant SAN cells were similar to those of WT cells in all experimental groups.

**Supplementary Video 1** Representative video showing intracellular cAMP application (100 µM) to a SAN pacemaker cell via the pipette solution in the whole-cell patch clamp configuration. The SAN cell was isolated from a transgenic mouse expressing the cAMP FRET sensor Epac1-camps and FRET was continuously monitored as the ratio between emission at 517-694 nm (YFP) and emission at 454-508 nm (CFP), upon excitation at 445 nm. The video visualizes the change in FRET ratio upon establishing the whole-cell configuration, indicating binding of cAMP to the sensor throughout the cell. Video plays at 10x speed. For details see methods and text in the main manuscript.

**Supplementary Video 2** Airy-scan 2 superresolution microscope movie of intracellular calcium fluorescence signals. Rhythmic and global calcium signals recorded in a WT whole-mount SAN preparation in control (before perfusion with Carbenoxolone).

**Supplementary Video 3** Airy-scan 2 superresolution microscope movie of intracellular calcium fluorescence signals recorded in a WT whole-mount SAN preparation (the same as in Supplementary Movie 2) after perfusion with Carbenoxolone (100 µM). The signal is slower and arrhythmic compared to the control (see Supplementary Movie 2).

**Supplementary Video 4** Airy-scan 2 superresolution microscope movie of intracellular calcium fluorescence signals. Rhythmic and global calcium signals recorded in a WT whole-mount SAN preparation in control (before perfusion with 1-Heptanol).

**Supplementary Video 5** Airy-scan 2 superresolution microscope movie of intracellular calcium fluorescence signals recorded in a WT whole-mount SAN preparation (the same as in Supplementary Movie 4) after perfusion with 1-Heptanol (10 µM). The signal is slower and arrhythmic compared to the control (see Supplementary Movie 4).

### Model supplement

#### 1 Model Description

The sinoatrial node is modeled as a 3-dimensional cuboid of $N = 500$ locally coupled phase oscillators. Each oscillator is randomly assigned a coordinate tuple $\vec{n} = (n_x, n_y, n_z)$ with $1 \leq n_x \leq N_x = 60$, $1 \leq n_y \leq N_y = 12$, and $1 \leq n_z \leq N_z = 4$. With the boundaries $N_{x,y,z}$ we mimicked the spatial outline of the sinoatrial node. Each oscillator is symmetrically coupled to its 5 nearest neighbors according to the Euclidean distance of the index tuples. The subset of index tuples of the oscillators connected to oscillator $\vec{n} = (n_x, n_y, n_z)$ is denoted as $\mathcal{C}(\vec{n})$.

Oscillators are randomly assigned to the subpopulations "type 0" and "type 1" according to the experimentally observed fractions (reported in Methods).

The state of an oscillator $\vec{n}$ at time $t$ is described by its phase variable $\phi_{\vec{n}}(t) \in [0, 2\pi)$, which progresses according to the differential equation

$$\frac{\mathrm{d}}{\mathrm{d}t}\phi_{\vec{n}}(t) = \frac{2\pi}{T_{\vec{n}}} + \sigma \sum_{\vec{n}' \in \mathcal{C}(\vec{n})} \Phi_{\vec{n}}(\phi_{\vec{n}} - \phi_{\vec{n}'})\, \delta\left(t - \frac{\phi_{\vec{n}'}}{2\pi} T_{\vec{n}'}\right) + \xi_{\vec{n}}(t) \; . \tag{1}$$

Here $T_{\vec{n}}$ denotes the unperturbed period of the oscillator (see below), $\sigma$ is the coupling strength (a free parameter 1), $\Phi_{\vec{n}}$ is the phase response curve (PRC) of the oscillator and $\delta$ denotes the Dirac delta distribution indicating that a phase change only occurs when oscillator $\vec{n}'$ expresses an action potential ($\phi_{\vec{n}'} = 0$). Finally, $\xi_{\vec{n}}(t)$ models uncorrelated Gaussian noise with mean 0 and variance $(2\pi/\mathrm{SNR})^2/\mathrm{d}t$, where the signal to noise ratio SNR is a second free parameter. Also, note that the variance is corrected for the time binning $\mathrm{d}t$ of the simulation.

**Oscillation periods $T_{\vec{n}}$.** Oscillation periods $T_{\vec{n}}$ are randomly assigned to each oscillator according to Gaussian distributions with mean and variance matching those of experimentally sampled populations (reported in Methods). We thereby distinguish between empirically determined values for type 0 and type 1 oscillators.

**PRCs $\Phi_{\vec{n}}$.** The PRC $\Phi(\phi)$ for oscillator $\vec{n}$ used in the theory of phase coupled oscillators is related to the experimentally measured period $T'(t)$ with perturbation at time $t$ after an AP as

$$\Phi\left(\frac{2\pi t}{T_{\vec{n}}}\right) = 1 - \frac{T'(t)}{T_{\vec{n}}} \ . \tag{2}$$

PRCs $\Phi$ used in our siunulations are matched to experimental data in two steps. 1) We fitted model functions $F_{0/1}(\phi)$ to the mean experimentally obtained PRC. We thereby distinguished between cells with type 0 and type 1 PRCs and selected the model function according to the type of simulated oscillator type($\vec{n}$). For type 0 PRCs the model function was

$$F_0(\phi) = \begin{cases} \frac{-\vartheta_2}{(\phi-\vartheta_0)^2+\vartheta_1} + \frac{\vartheta_2}{\vartheta_0^2+\vartheta_1} & 0 \le \vartheta_0 - \phi < \pi \\ \frac{\vartheta_3}{(\phi-\vartheta_0)^2+\vartheta_4} - \frac{\vartheta_3}{(2\pi-\vartheta_0)^2+\vartheta_4} & -\pi \le \vartheta_0 - \phi < 0 \end{cases} \tag{3}$$

For type 1 PRCs the model function was

$$F_1(\phi) = \vartheta_2 \, M(\phi, \vartheta_0, \vartheta_1) - \vartheta_5 \, M(\phi, \vartheta_3, \vartheta_4) \tag{4}$$

with

$$M(\phi, \mu, \kappa) = \exp[\kappa\,(\cos(\phi-\mu) - 1)] - \exp[\kappa\,(\cos(\mu) - 1)] \ .$$

The parameters $\vartheta_i$ were used for fitting.

2) To account for a finite action potential width that is larger than the pulses used to probe the PRCs in the in-vitro experiments we obtained the effective PRCs $\Phi_{\vec{n}}(\phi)$ by convolving the fitted model functions $F_{\text{type } 0,1}(\phi)$ with a stylized action potential shape $A(\phi)$, i.e.,

$$\Phi_{\vec{n}}(\phi) = \int_0^{2\pi} \mathrm{d}\varphi A_{\vec{n}}(\varphi) F_{\text{type}(\vec{n})}(\phi - \varphi) \ . \tag{5}$$

To restrict free parameters to a minimum, the shape of the action potential is assumed triangular

$$A_{\vec{n}}(\phi) = \max\left(0, 1 - \frac{\phi}{2\pi}\frac{T_{\vec{n}}}{2\,wT_0}\right)\frac{T_{\vec{n}}}{2\pi\,wT_0}$$

with half width $w$ in units of the mean unperturbed period $T_0$ of type 0 oscillators. Note that the action potential shape as a function of phase $\phi$ was adjusted to the period length $T_{\vec{n}}$ of the oscillator in order to result in a constant width $wT_0$ in time that is independent of the oscillator. The AP width $w$ is the third free parameter. Furthermore, the amplitude of the AP is normalized such that the integral over the PRC does not

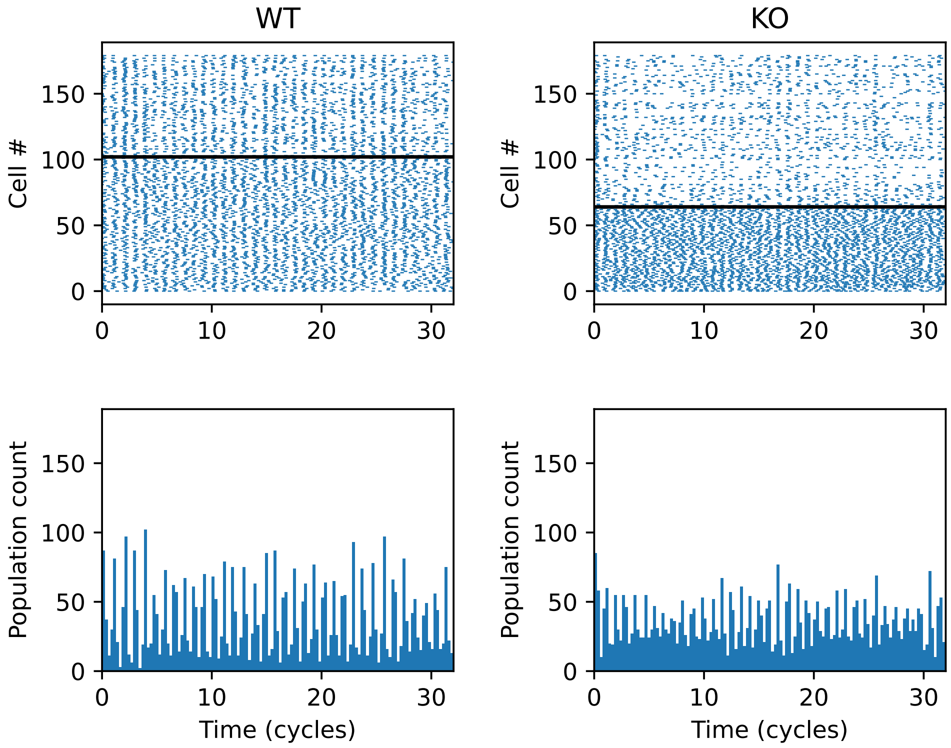


**Supplementary Figure 4**: Simulations results with large difference in synchronization ($\sigma = 0.3, w = 0.19$, SNR$=3$). Top: Spike raster plots. Every dots indicates an AP. Solid black lines indicates the boundary between type 1 (below black line) and type 0 (above black line) oscillators. Bottom: population count. Number of APs per phase bin (1/4 cycle relative to $T_0$). Left and right columns correspond to simulations obtained with PRCs fitted to recordings from WT and HCN1 KO cells.

change, i.e.,

$$\int_0^{2\pi} \mathrm{d}\varphi \, A(\varphi) = 1 \ .$$

#### 2 Model behavior

Depending on the choice of free parameters, SNR, AP width $w$ and coupling strength $\sigma$, simulations can exhibit large differences in synchronization. In the example shown in Fig. **4**, entrainment and jitter is much higher in simulations with PRCs fitted to wild type (WT) recordings as compared to recordings obtaine in tissue from HCN1 KO mice

To more systematically explore the parameter dependence we assessed simulations by means of three quality measures. The mean period, the jitter (variability of the period as determined from the mean autocorrelation), and, most importantly, the entrainment $E$. The entrainment measures the fraction of oscillators that are active in each population cycle and therefore assumes values between 0 and 1. Parameter dependence of mean period and period jitter did not exhibit strong differences between WT and HNC1 KO group (Fig. **5**A), whereas entrainment was much worse in the HCN1KO group for physiologically reasonable broad APs, low coupling strength, and intermediate SNR (Fig. **5**B, Left). Comparing the entrainment of the HCN4FEA with and without 100 $\mu$M cAMP with their respective controls did not produce such clear differences.

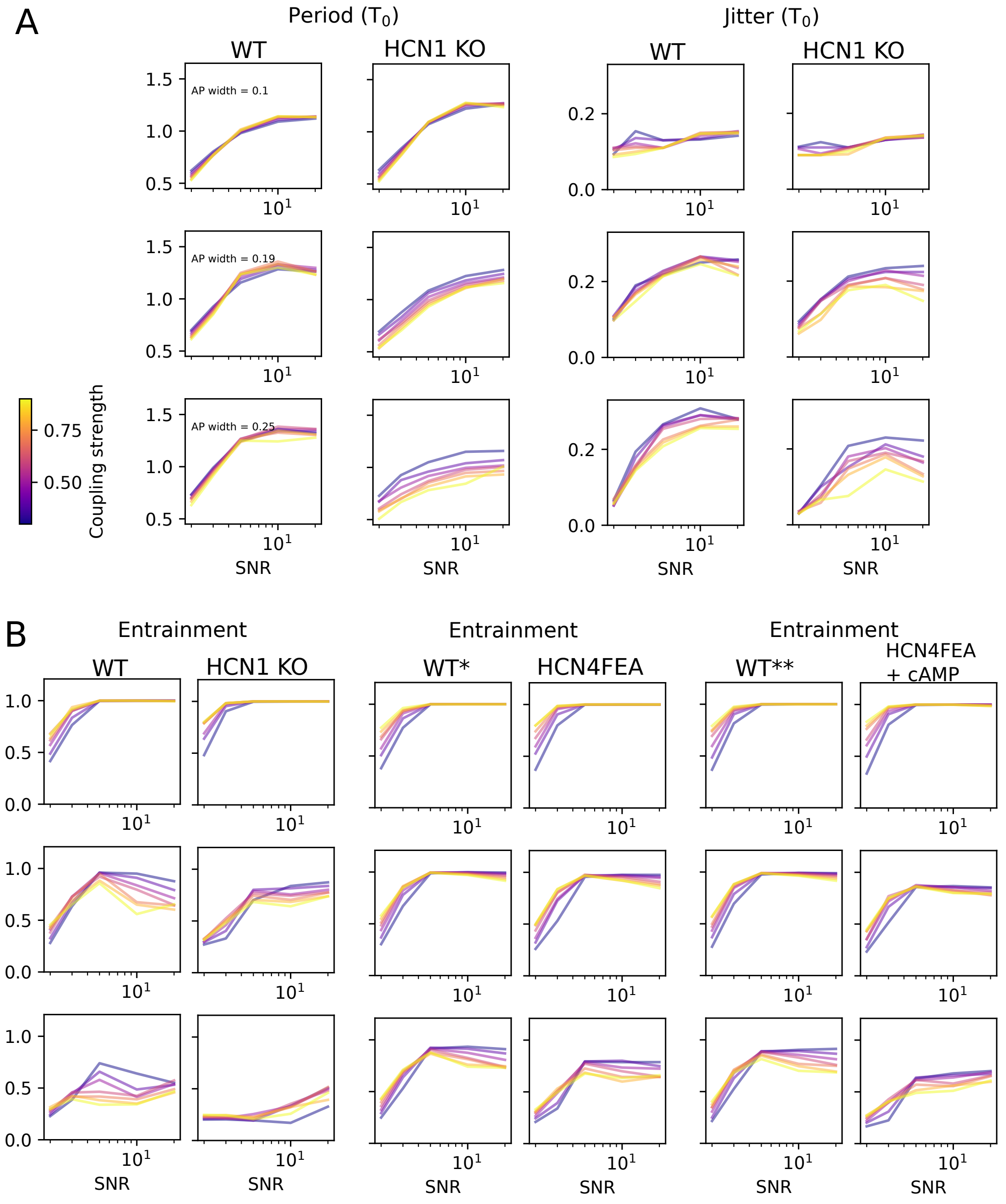


**Supplementary Figure 5**: Parameter scans. (A) Mean period of an oscillator and jitter around mean period increase with SNR almost independently of AP width $w$, and only show mild dependence on coupling strength $\sigma$ (color code). Most importantly, only small differences are observable between WT and experimental group except for a general decrease of mean period length for longer AP widths. (B) Entrainment (average participation per cycle) for all three experimental comparison groups exhibits differences in the HCN1KO experimental group but not for HCN4FEA and HCN4FEA + cAMP groups.

#### Supplementary References


1. Fenske S, Hennis K, Rotzer RD, Brox VF, Becirovic E, Scharr A, Gruner C, Ziegler T, Mehlfeld V, Brennan J, Efimov IR, Pauza AG, Moser M, Wotjak CT, Kupatt C, Gonner R, Zhang R, Zhang H, Zong X, Biel M, Wahl-Schott C. Camp-dependent regulation of hcn4 controls the tonic entrainment process in sinoatrial node pacemaker cells. *Nat Commun*. 2020;11:5555
2. Fenske S, Krause SC, Hassan SI, Becirovic E, Auer F, Bernard R, Kupatt C, Lange P, Ziegler T, Wotjak CT, Zhang H, Hammelmann V, Paparizos C, Biel M, Wahl-Schott CA. Sick sinus syndrome in hcn1-deficient mice. *Circulation*. 2013;128:2585-2594
3. Hennis K, Rotzer RD, Rilling J, Wu Y, Thalhammer SB, Biel M, Wahl-Schott C, Fenske S. In vivo and ex vivo electrophysiological study of the mouse heart to characterize the cardiac conduction system, including atrial and ventricular vulnerability. *Nat Protoc*. 2022;17:1189-1222
4. Moghtadaei M, Dorey TW, Rose RA. Evaluation of non-linear heart rate variability using multi-scale multi-fractal detrended fluctuation analysis in mice: Roles of the autonomic nervous system and sinoatrial node. *Front Physiol*. 2022;13:970393
5. Usai DS, Aasum E, Thomsen MB. The isolated, perfused working heart preparation of the mouse-advantages and pitfalls. *Acta Physiol (Oxf)*. 2025;241:e70023
6. El Khoury N, Mathieu S, Marger L, Ross J, El Gebeily G, Ethier N, Fiset C. Upregulation of the hyperpolarization-activated current increases pacemaker activity of the sinoatrial node and heart rate during pregnancy in mice. *Circulation*. 2013;127:2009-2020
7. Shemla O, Tsutsui K, Behar JA, Yaniv Y. Beating rate variability of isolated mammal sinoatrial node tissue: Insight into its contribution to heart rate variability. *Front Neurosci*. 2020;14:614141
8. Yaniv Y, Ahmet I, Tsutsui K, Behar J, Moen JM, Okamoto Y, Guiriba TR, Liu J, Bychkov R, Lakatta EG. Deterioration of autonomic neuronal receptor signaling and mechanisms intrinsic to heart pacemaker cells contribute to age-associated alterations in heart rate variability in vivo. *Aging Cell*. 2016;15:716-724
9. Hoesl E, Stieber J, Herrmann S, Feil S, Tybl E, Hofmann F, Feil R, Ludwig A. Tamoxifen-inducible gene deletion in the cardiac conduction system. *J Mol Cell Cardiol*. 2008;45:62-69
10. Baruscotti M, Bucchi A, Viscomi C, Mandelli G, Consalez G, Gnecchi-Rusconi T, Montano N, Casali KR, Micheloni S, Barbuti A, DiFrancesco D. Deep bradycardia and heart block caused by inducible cardiac-specific knockout of the pacemaker channel gene hcn4. *Proceedings of the National Academy of Sciences of the United States of America*. 2011;108:1705-1710
11. Mesirca P, Alig J, Torrente AG, Muller JC, Marger L, Rollin A, Marquilly C, Vincent A, Dubel S, Bidaud I, Fernandez A, Seniuk A, Engeland B, Singh J, Miquerol L, Ehmke H, Eschenhagen T, Nargeot J, Wickman K, Isbrandt D, Mangoni ME. Cardiac arrhythmia induced by genetic silencing of 'funny' (f) channels is rescued by girk4 inactivation. *Nat Commun*. 2014;5:4664
12. Kozasa Y, Nakashima N, Ito M, Ishikawa T, Kimoto H, Ushijima K, Makita N, Takano M. Hcn4 pacemaker channels attenuate the parasympathetic response and stabilize the spontaneous firing of the sinoatrial node. *J Physiol*. 2018;596:809-825
13. Harzheim D, Pfeiffer KH, Fabritz L, Kremmer E, Buch T, Waisman A, Kirchhof P, Kaupp UB, Seifert R. Cardiac pacemaker function of hcn4 channels in mice is confined to embryonic development and requires cyclic amp. *EMBO J*. 2008;27:692-703
14. Alig J, Marger L, Mesirca P, Ehmke H, Mangoni ME, Isbrandt D. Control of heart rate by camp sensitivity of hcn channels. *Proceedings of the National Academy of Sciences of the United States of America*. 2009;106:12189-12194
15. Ren L, Thai PN, Gopireddy RR, Timofeyev V, Ledford HA, Woltz RL, Park S, Puglisi JL, Moreno CM, Santana LF, Conti AC, Kotlikoff MI, Xiang YK, Yarov-Yarovoy V, Zaccolo M, Zhang XD, Yamoah EN, Navedo MF, Chiamvimonvat N. Adenylyl cyclase isoform 1 contributes to sinoatrial node automaticity via functional microdomains. *JCI Insight*. 2022;7
16. Piantoni C, Carnevali L, Molla D, Barbuti A, DiFrancesco D, Bucchi A, Baruscotti M. Age-related changes in cardiac autonomic modulation and heart rate variability in mice. *Front Neurosci*. 2021;15:617698
17. Landi S, Giannetti F, Benzoni P, Campostrini G, Rossi G, Piantoni C, Bertoli G, Bonfanti C, Carnevali L, Bucchi A, Baruscotti M, Careccia G, Messina G, Barbuti A. Lack of the transcription factor nfix causes tachycardia in mice sinus node and rats neonatal cardiomyocytes. *Acta Physiol (Oxf)*. 2023;239:e13981

18. Lakin R, Guzman C, Izaddoustdar F, Polidovitch N, Goodman JM, Backx PH. Changes in heart rate and its regulation by the autonomic nervous system do not differ between forced and voluntary exercise in mice. *Front Physiol*. 2018;9:841
19. Uechi M, Asai K, Osaka M, Smith A, Sato N, Wagner TE, Ishikawa Y, Hayakawa H, Vatner DE, Shannon RP, Homcy CJ, Vatner SF. Depressed heart rate variability and arterial baroreflex in conscious transgenic mice with overexpression of cardiac gsalpha. *Circ Res*. 1998;82:416-423
20. Rose RA, Kabir MG, Backx PH. Altered heart rate and sinoatrial node function in mice lacking the camp regulator phosphoinositide 3-kinase-gamma. *Circ Res*. 2007;101:1274-1282
21. Mesquita T, Zhang R, Cho JH, Zhang R, Lin YN, Sanchez L, Goldhaber JI, Yu JK, Liang JA, Liu W, Trayanova NA, Cingolani E. Mechanisms of sinoatrial node dysfunction in heart failure with preserved ejection fraction. *Circulation*. 2022;145:45-60
22. Herrmann S, Stieber J, Stockl G, Hofmann F, Ludwig A. Hcn4 provides a 'depolarization reserve' and is not required for heart rate acceleration in mice. *EMBO J*. 2007;26:4423-4432
23. Lolicato M, Bucchi A, Arrigoni C, Zucca S, Nardini M, Schroeder I, Simmons K, Aquila M, DiFrancesco D, Bolognesi M, Schwede F, Kashin D, Fishwick CW, Johnson AP, Thiel G, Moroni A. Cyclic dinucleotides bind the c-linker of hcn4 to control channel camp responsiveness. *Nat Chem Biol*. 2014;10:457-462
24. Wu Y, Wang Q, Granger J, Reyes Gaido O, Lopez-Cecetaite G, Aguilar EN, Ludwig A, Moroni A, Bianchet MA, Anderson ME. Hcn4 channels sense temperature and determine heart rate responses to heat. *Nat Commun*. 2025;16:2102
25. Dorey TW, McRae MD, Belke DD, Rose RA. Pde4d mediates impaired beta-adrenergic receptor signalling in the sinoatrial node in mice with hypertensive heart disease. *Cardiovasc Res*. 2023;119:2697-2711
26. Gao Z, Rasmussen TP, Li Y, Kutschke W, Koval OM, Wu Y, Wu Y, Hall DD, Joiner ML, Wu XQ, Swaminathan PD, Purohit A, Zimmerman K, Weiss RM, Philipson KD, Song LS, Hund TJ, Anderson ME. Genetic inhibition of na+-ca2+ exchanger current disables fight or flight sinoatrial node activity without affecting resting heart rate. *Circ Res*. 2013;112:309-317
27. D'Souza A, Bucchi A, Johnsen AB, Logantha SJ, Monfredi O, Yanni J, Prehar S, Hart G, Cartwright E, Wisloff U, Dobryznski H, DiFrancesco D, Morris GM, Boyett MR. Exercise training reduces resting heart rate via downregulation of the funny channel hcn4. *Nat Commun*. 2014;5:3775
28. D'Souza A, Wang Y, Anderson C, Bucchi A, Baruscotti M, Olieslagers S, Mesirca P, Johnsen AB, Mastitskaya S, Ni H, Zhang Y, Black N, Cox C, Wegner S, Bano-Otalora B, Petit C, Gill E, Logantha S, Dobrzynski H, Ashton N, Hart G, Zhang R, Zhang H, Cartwright EJ, Wisloff U, Mangoni ME, da Costa Martins PA, Piggins HD, DiFrancesco D, Boyett MR. A circadian clock in the sinus node mediates day-night rhythms in hcn4 and heart rate. *Heart Rhythm*. 2021;18:801-810
29. Neco P, Torrente AG, Mesirca P, Zorio E, Liu N, Priori SG, Napolitano C, Richard S, Benitah JP, Mangoni ME, Gomez AM. Paradoxical effect of increased diastolic ca(2+) release and decreased sinoatrial node activity in a mouse model of catecholaminergic polymorphic ventricular tachycardia. *Circulation*. 2012;126:392-401
